# Fluorescence lifetime-based FRET biosensors for monitoring N-terminal domain interactions of TDP-43 in living cells: A novel resource for ALS and FTD drug discovery

**DOI:** 10.1101/2024.10.11.617905

**Authors:** Noah Nathan Kochen, Marguerite Murray, Nagamani Vunnam, Elly E. Liao, Lihsia Chen, Anthony R. Braun, Jonathan N. Sachs

**Author notes:** First author (Noah Nathan Kochen).

## Abstract

TAR DNA-binding protein 43 (TDP-43) pathological aggregates are widely implicated in Alzheimer’s disease, frontotemporal dementia and amyotrophic lateral sclerosis. While therapeutic platforms targeting TDP-43 have predominantly targeted its aggregation, recent findings suggest that loss of functional TDP-43 dimers and multimers — essential for RNA processing — occur upstream of aggregation and is driven through disruption of N-terminal domain (NTD) interactions. Here, we demonstrate that these interactions are targetable via cellular fluorescence lifetime-based FRET biosensors which we used to screen the FDA-approved Selleck library. Our NTD-specific hit ketoconazole rescues sorbitol-induced TDP-43 mislocalization and aggregation, and ameliorates TDP-43 induced downregulation of SREBP2, a TDP-43 mRNA binding target with known implication in ALS. In addition, ketoconazole improves neurite outgrowth in a TDP-43 overexpressing neuron model and motor dysfunction in TDP-43 overexpressing C. elegans. Taken together, our platform represents a novel approach for targeting NTD-dependent TDP-43 interactions, and the identification of ketoconazole validates an exciting translational premise for TDP-43 drug discovery.

## Introduction

TAR DNA-binding protein 43 (TDP-43) proteinopathies are present in roughly 75% of Alzheimer’s disease (AD), 97% of amyotrophic lateral sclerosis (ALS) and 50% of frontotemporal dementia (FTD) cases, spanning both sporadic and familial forms of these diseases^1^. The pathophysiology of these diseases spans a series of TDP-43 gain of toxic functions (e.g., neuroinflammation, mitochondrial dysfunction and proteostatic stress driven by mislocalization and aggregation of TDP-43) and loss of native functions (e.g., loss of RNA-processing, reversible stress granule formation and axonal transport)^2,3^. Due to the widespread presence of TDP-43 cytoplasmic inclusions in late-stage ALS and FTD^4^, numerous small molecule screening strategies have focused on inhibiting its pathological aggregation^5-7^.

TDP-43 is a multi-domain protein involved in RNA stability, splicing and transport as well as formation of cytosolic stress granules during cell stress^8^. TDP-43 contains a globular N-terminal domain (NTD) involved in its native self-oligomerization, two globular RNA recognition motifs (RRMs) and a glycine-rich disordered C-terminal domain (CTD) implicated in the formation of amyloid aggregates^9^. Assemblies formed via CTD-CTD interactions are important for formation of cytoplasmic stress granules in response to cellular stress; however, if unresolved, these assemblies can lead to irreversible aggregation linked to TDP-43’s toxic gain of function pathology^10,11^.

However, not all TDP-43 assemblies are detrimental to cellular health. TDP-43’s nuclear retention and native functions — RNA-binding and splicing activity — require oligomerization mediated by NTD-NTD interactions^12^. Furthermore, eliminating NTD-mediated oligomers via point mutation renders TDP-43 more susceptible to mislocalization, aggregation, and reduces its resident cellular half-life^13^. A recent study by Oiwa et al. identified a significant loss of functional NTD-mediated TDP-43 oligomers in the brains and spinal cords of ALS patients^14^. This pathological decrease in TDP-43 multimerization *<u>precedes</u>* toxic gain of function, namely TDP-43 mislocalization to the cytoplasm, subsequent unresolved aggregation, and hyperphosphorylation^15-28^.

This study builds upon those findings through the development of a novel therapeutic discovery platform that monitors NTD-dependent interactions of functional TDP-43 multimers in living cells. Hit compounds identified via our platform confirm the hypothesis that small molecules that stabilize NTD-dependent self-assembly of functional multimeric TDP-43 are capable of rescuing TDP-43 associated pathological phenotypes. We have engineered a series of live-cell, fluorescence lifetime (FLT) based Förster resonance energy transfer (FRET) biosensors that monitor TDP-43 homo/self-assemblies while discriminating between NTD-dependent and NTD-independent interactions. Our previous work employed a similar FLT-FRET strategy for high-throughput screening (HTS) campaigns targeting other neurodegenerative disease associated proteins (e.g., alpha-synuclein, tau, and huntingtin) which, similar to TDP-43, can populate an ensemble of aggregation and native conformation states^29-32^.

With our TDP-43 biosensors, we screened the 2,684-compound FDA-approved Selleck library and identified ketoconazole, a known P450 cytochrome inhibitor that increases TDP-43 FRET in an NTD-dependent manner. The design of our biosensors and use of counter-screens allows us to identify hit compounds that bias the TDP-43 ensemble toward a specific distribution — i.e., stabilizing NTD-interaction dependent nuclear assemblies. The identification of NTD-dependent small molecules capable of rescuing TDP-43-induced deficits demonstrates that our FLT-FRET biosensor platform represents a novel and exciting therapeutic discovery strategy for FTD, ALS, and other TDP-43 proteinopathies.

## Results

### Intermolecular FRET biosensor monitors NTD-dependent TDP-43 interactions

Biosensor constructs were engineered using a C-terminal fusion of either donor (mNeonGreen, mNg) or acceptor (mCherry, mCh) fluorescent proteins to full-length (FL) or NTD deletion (ΔNTD) human TDP-43 to generate FL TDP-43-mNg, FL TDP-43-mCh, and ΔNTD TDP-43-mCh (Figure 1A). The mNg/mCh FRET pair provides an improvement over our previously GFP/RFP biosensors due to the fluorophores’ high quantum yields and robust spectral overlap^33^. In addition to TDP-43 biosensors, donor-only mNg and donor-acceptor mCh-Linker-mNg ((GGGGS)_14_ linker) biosensors were used as controls.

**Figure 1.**
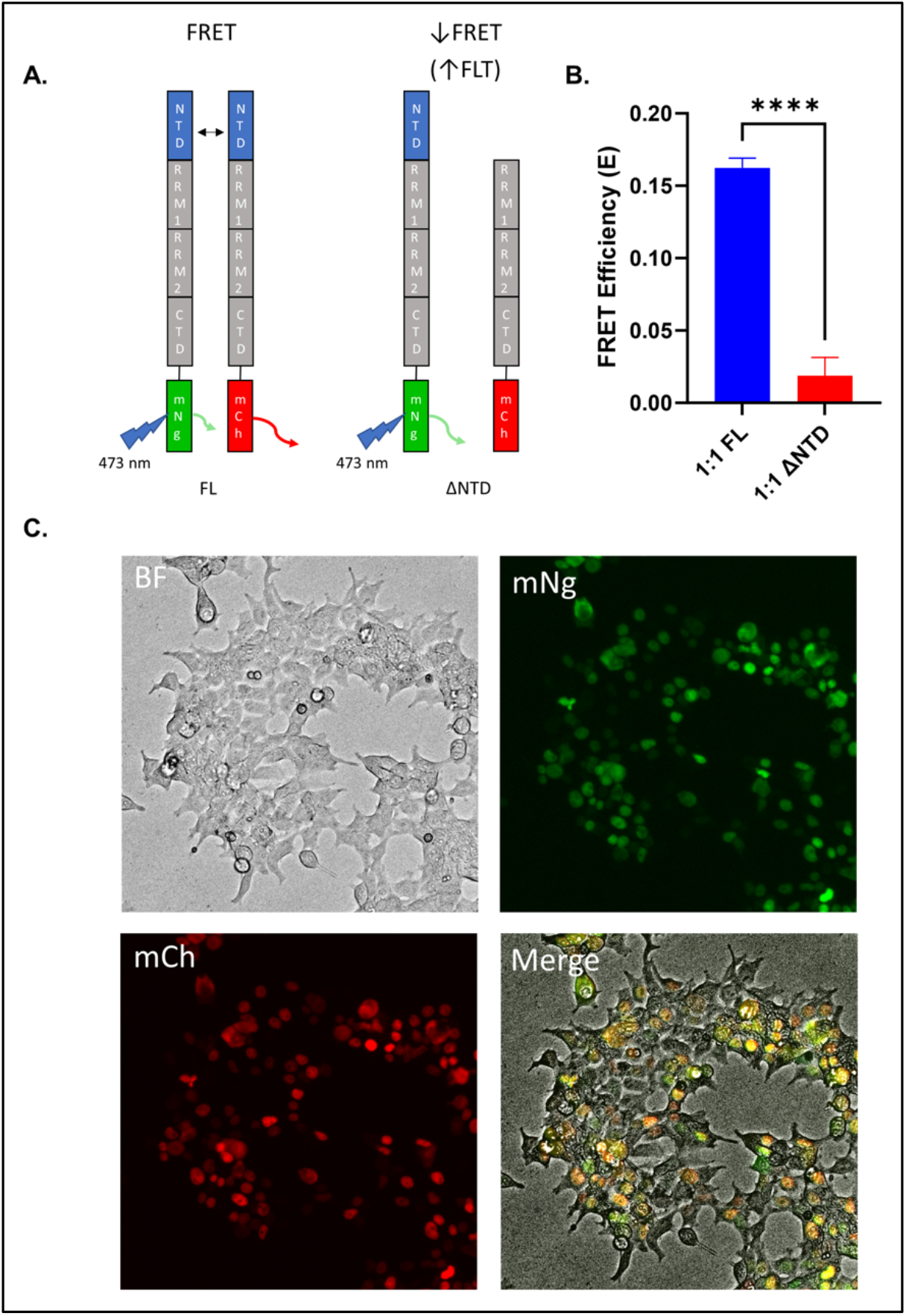
FLT-FRET TDP-43 biosensor monitors nuclear NTD-dependent TDP-43 interactions. **(A)** Diagrams of full-length (WT) and NTD-null (ΔNTD) TDP-43-fluorophore fusion biosensors. **(B)** FRET of FL and ΔNTD biosensors under 1:1 donor:acceptor ratio in HEK293T cells after 24 hours of expression. **(C)** Fluorescence live-cell imaging of 1:1 FL biosensor expressed for 24 hours in HEK293T cells. Statistic shown is a one-way ANOVA comparison with Bonferroni correction (subset of full donor:acceptor ratio experiment shown in Supplemental Figure 2, ****p < 0.0001).

Biosensor FRET is measured by monitoring the donor’s FLT with (*τ*_DA_) and without (*τ*_D_) expression of the acceptor construct, where FRET efficiency is E=1-*τ*_DA_/*τ*_D_. Because FLT is an intrinsic property of the fluorophore, it provides increased sensitivity relative to intensity-based approaches^34^. Supplemental Figure 1 illustrates the ∼16-fold reduction in variability for FLT versus fluorescence intensity — which allows for robust compound detection — for our TDP-43 and linker biosensors.

Figure 1B shows that expression of the FL TDP-43-mNg + FL TDP-43-mCh at 1:1 ratio results in a significantly higher FRET efficiency (E = 0.16) compared to the FL TDP-43-mNg + ΔNTD TDP-43-mCh biosensor (E = 0.02). The FRET efficiency from the ΔNTD system is at similar levels observed for non-interacting background FRET biosensors used in our previous studies^29,30^. Interestingly, when we hold the amount of donor construct constant and add increasing acceptor — increasing the total amount of TDP-43 — we observe a concomitant increase in FRET for both the FL and ΔNTD biosensors, suggesting the ensemble of TDP-43 assemblies is being biased toward non-NTD dependent species at higher levels of protein expression (Supplemental Figure 2).

Although increasing the total amount of TDP-43 results in elevated FL FRET (e.g., a stronger FRET signal), the concomitant increase in NTD-independent FRET is problematic for specifically targeting NTD-dependent interactions; hence, all subsequent TDP-43 FRET assays are performed using the 1:1 biosensor condition. Figure 1C shows that expression of the FL biosensors at this ratio results in diffuse nuclear localization of TDP-43, devoid of puncta or cytoplasmic mislocalization. Biosensor expression was confirmed via immunoblot assay of biosensor expressing HEK293T cells (Supplemental Figure 3).

We next confirmed that the ΔNTD biosensor resulted in a loss of TDP-43 multimers using a disuccinimidyl glutarate (DSG) cross-linking protocol in HEK293T cells expressing endogenous TDP-43 or transiently transfected with FL, or ΔNTD TDP-43-mNg constructs. As expected, DSG cross-linking resulted in dimeric and multimeric TDP-43 bands across all conditions due to endogenous TDP-43 expression (Supplemental Figure 4A). The addition of FL TDP-43-mNg resulted in the highest amount of cross-linked TDP-43 dimer/multimers, whereas ΔNTD expressing conditions only shows the endogenous TDP-43 dimer/multimers. Densitometry analysis of the multimeric TDP-43 bands relative to total protein staining (Supplemental Figure 4B-C and Supplemental Figure 5) confirms these observations. Taken together, our data suggests that our biosensors report on nuclear NTD-dependent TDP-43 interactions in living cells.

### Selleck library FLT-FRET HTS

Prior to implementing our HTS pipeline we determined each biosensors coefficient of variation (CV) in the 1536-well plate format. Figure 2A-B confirms a robust FLT and FRET signal. It is important to note that in determining FRET there is error propagation due to the two separate FLT measurements (i.e., donor only and donor + acceptor), such that for all HTS applications, hit compounds are identified via FLT change relative to DMSO-treated wells to improve the sensitivity of the screen.

**Figure 2.**
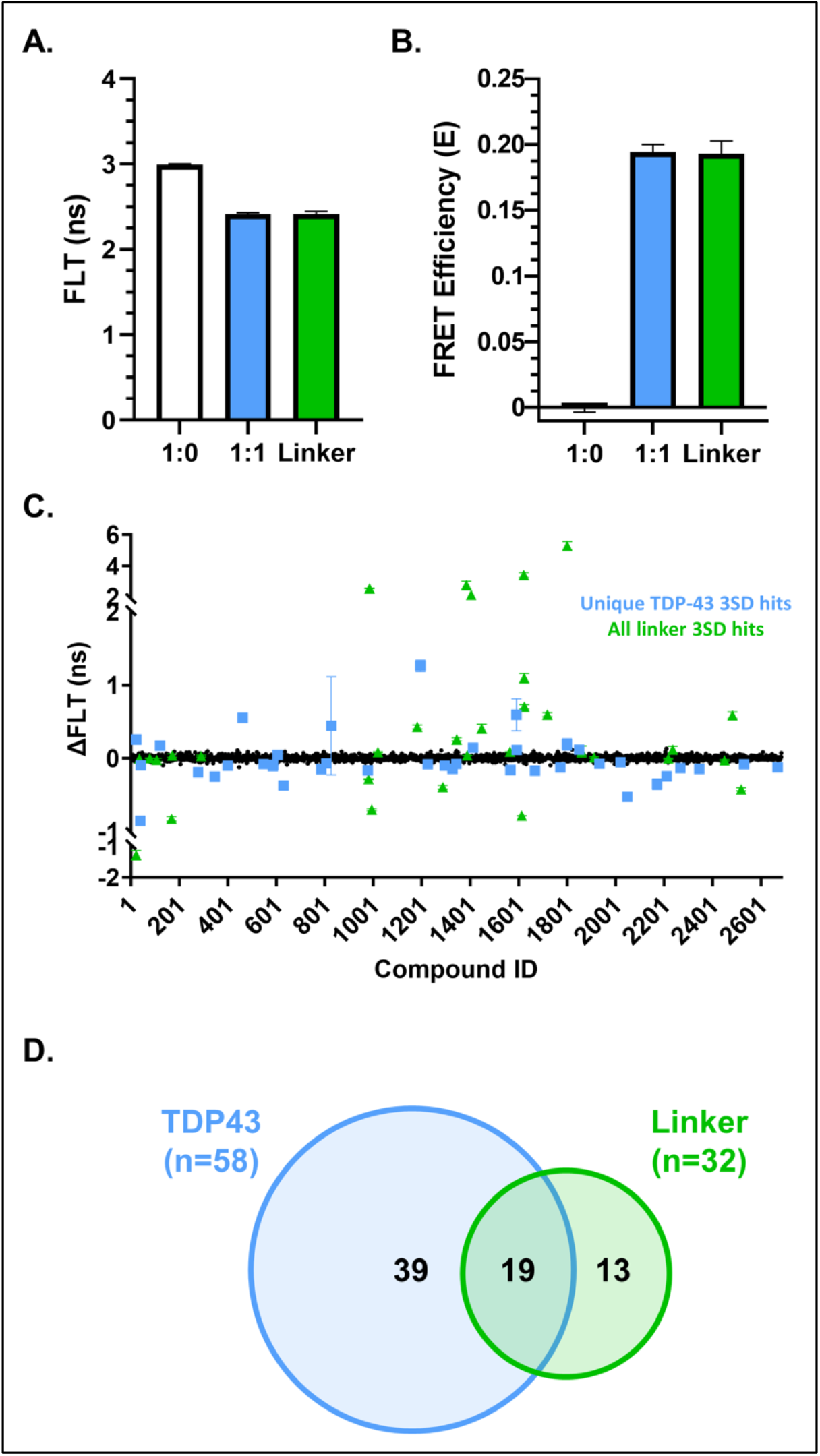
FDA-approved Selleck library screen in WT TDP-43 and linker control biosensors. **(A)** Average FLT of biosensors in HEK293T cells measured in 1536-well plate format for drug screening. **(B)** Average FRET efficiencies calculated from (A). **(C)** Average ΔFLT (FLT_drug_ – FLT_DMSO_) for FDA-approved Selleck library screened against FL and linker biosensors. Positive ΔFLT indicate molecules that decrease FRET, whereas negative ΔFLT indicate molecules that increase FRET. **(D)** Summary of TDP-43 and linker unique hits. Data shown are mean ± SEM from N=3 independent experiments.

Next, we executed our HTS of the FDA-approved, 2,684-compound Selleck library. Biosensor cells were plated into 1536-well drug plates with compounds at a final concentration of 10 μM. Figure 2C illustrates the summary data from triplicate screens using our FL TDP-43 biosensor and counter-screens using our linker-only control (diagrams of biosensors used in the HTS campaign shown in Supplemental Figure 6). The implementation of this linker counter-screen is essential to flag non-TDP-43 specific hits (e.g., those that non-specifically affect the XFPs). Compounds with a reproducible change in FLT (ΔFLT) of ±3 standard deviation (SD) across the triplicate screens that were not flagged in either the linker screen or the fluorescent interference spectral similarity filter^30^ were considered FL TDP-43 unique hits. Figure 2C-D shows TDP-43 screen summary data as change in fluorescent lifetime (ΔFLT) relative to untreated cells, highlighting ±3SD TDP-43 unique hits in light blue and molecules that were flagged as ±3SD hits in the control linker screens in green. Overall, we identified 58 hits, of which 19 were flagged in the linker counter-screen. Of the 39 unique TDP-43 hits, we excluded 16 due to known mechanism of action (MOA) associated with cell death and toxicity, resulting in 23 lead compounds (summarized in Supplemental Table 1).

### Identification of NTD-specific hit compounds via ΔNTD biosensor counter-screening

Next, using our ΔNTD biosensor, we implemented a counter-screen strategy to identify compounds that increase NTD-dependent interactions. An NTD-dependent compound will have a unique ΔFLT profile, specifically decreasing FLT or ΔFLT (increasing FRET) for the FL biosensor without affecting the ΔNTD biosensor. A summary of the counter-screen of the 23 TDP-43 unique hits at 10 *μ*M against the ΔNTD biosensor is reported in Supplemental Figure 7. Hits were flagged as NTD-dependent if FL and not ΔNTD ΔFLT showed a statistically significant difference relative to their respective biosensor’s DMSO condition. From these 23 compounds, ketoconazole emerged as the sole hit that reduced TDP-43 biosensor FLT and satisfied the FL vs ΔNTD hit flag (Figure 3A). Figure 3B illustrates a 7-point dose-response ketoconazole resulted in a FLT-FRET EC_50_ of 1.2 μM. Although not strictly meeting our NTD-dependent hit criterion, there are additional compounds that displayed a similar trend that are potentially worth further exploration. A discussion of these compounds is presented later in the manuscript.

**Figure 3.**
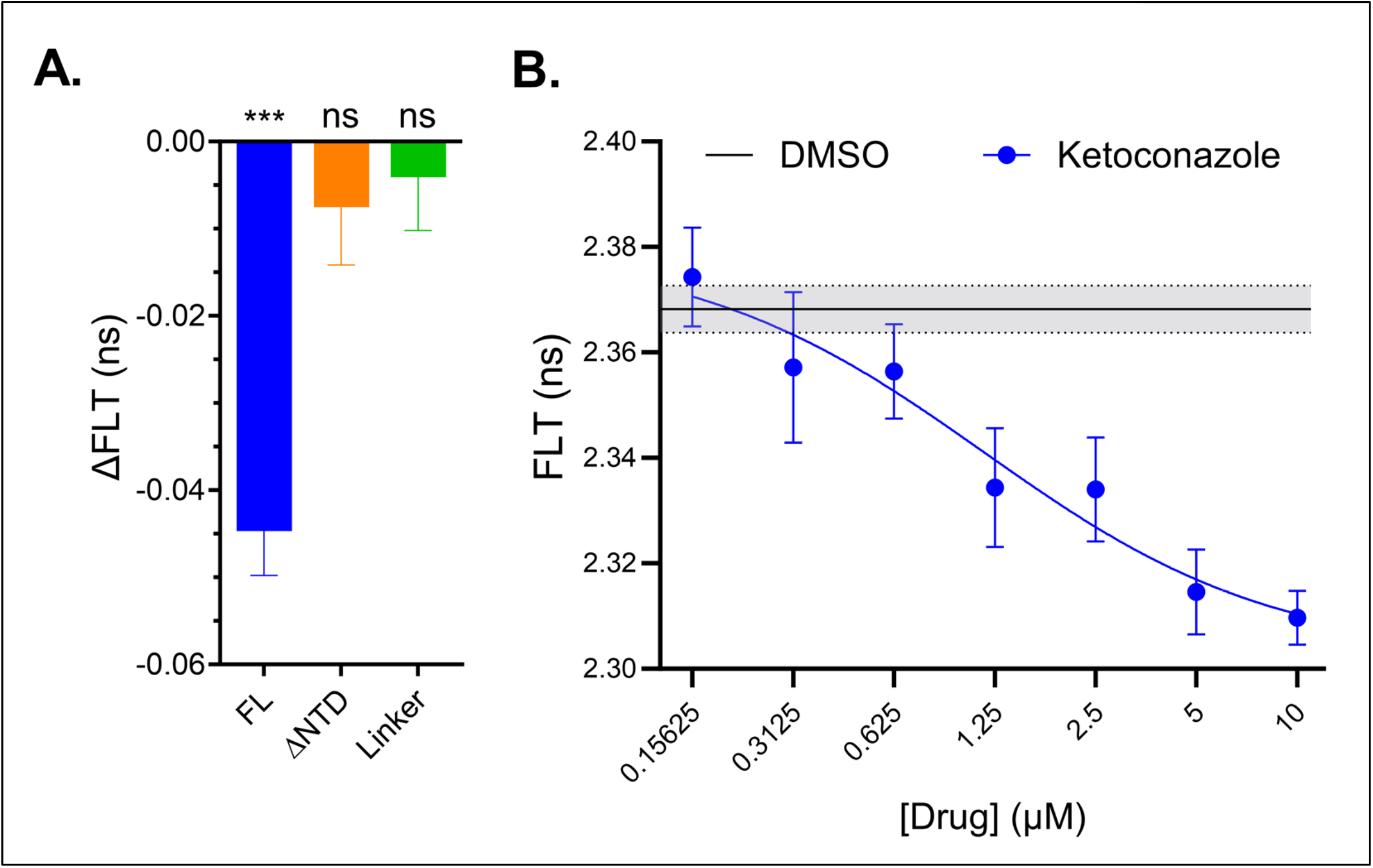
Ketoconazole modulates TDP-43 FLT in an NTD and dose dependent manner. **(A)** ΔFLT for ketoconazole at 10 μM (screening condition) tested against FL, ΔNTD and linker biosensors. **(B)** FLT dose response for FL biosensor treated with ketoconazole 7-point titration resulted in IC_50_ = 1.2μM. Statistics shown in (A) are one sample T tests to hypothetical mean of zero (DMSO baseline, ***p < 0.001). Data shown are mean ± SEM from N=3-6 independent experiments.

To ensure that the decrease in FLT induced by ketoconazole is not associated with TDP-43 aggregation and puncta formation, we imaged TDP-43-mNg expressing HEK293T cells treated with 10 *μ*M drug for 2 hours (the same treatment conditions for the HTS). Supplemental Figure 8A-B shows that ketoconazole does not induce TDP-43 aggregation, consistent with its NTD-dependent effect. In contrast, when we tested erdafitinib — a TDP-43 hit that increased FRET in the ΔNTD biosensor — we observed widespread formation of puncta and mislocalization of TDP-43 (Supplemental Figure 8C-D).

### Ketoconazole rescues loss of neurite outgrowth in TDP-43 overexpression model

Next, we tested ketoconazole’s ability to rescue TDP-43 induced neurite loss in a model of TDP-43 proteinopathy using differentiated Neuro2a (N2a) cells stably overexpressing FL TDP-43-mNg. Neurite outgrowth has been recently used as a robust phenotypic assay that monitors ALS associated cellular dysfunction and the discovery of novel ALS therapeutics^6^. FL TDP-43-mNg expression reduced neurite outgrowth by roughly 50% compared to naïve N2a cells (Figure 4A-B). Treatment of TDP-43-mNg expressing cells with 10 μM ketoconazole restored neurite outgrowth to 75% of naïve N2a cells, with DMSO-controls having no significant effect.

**Figure 4.**
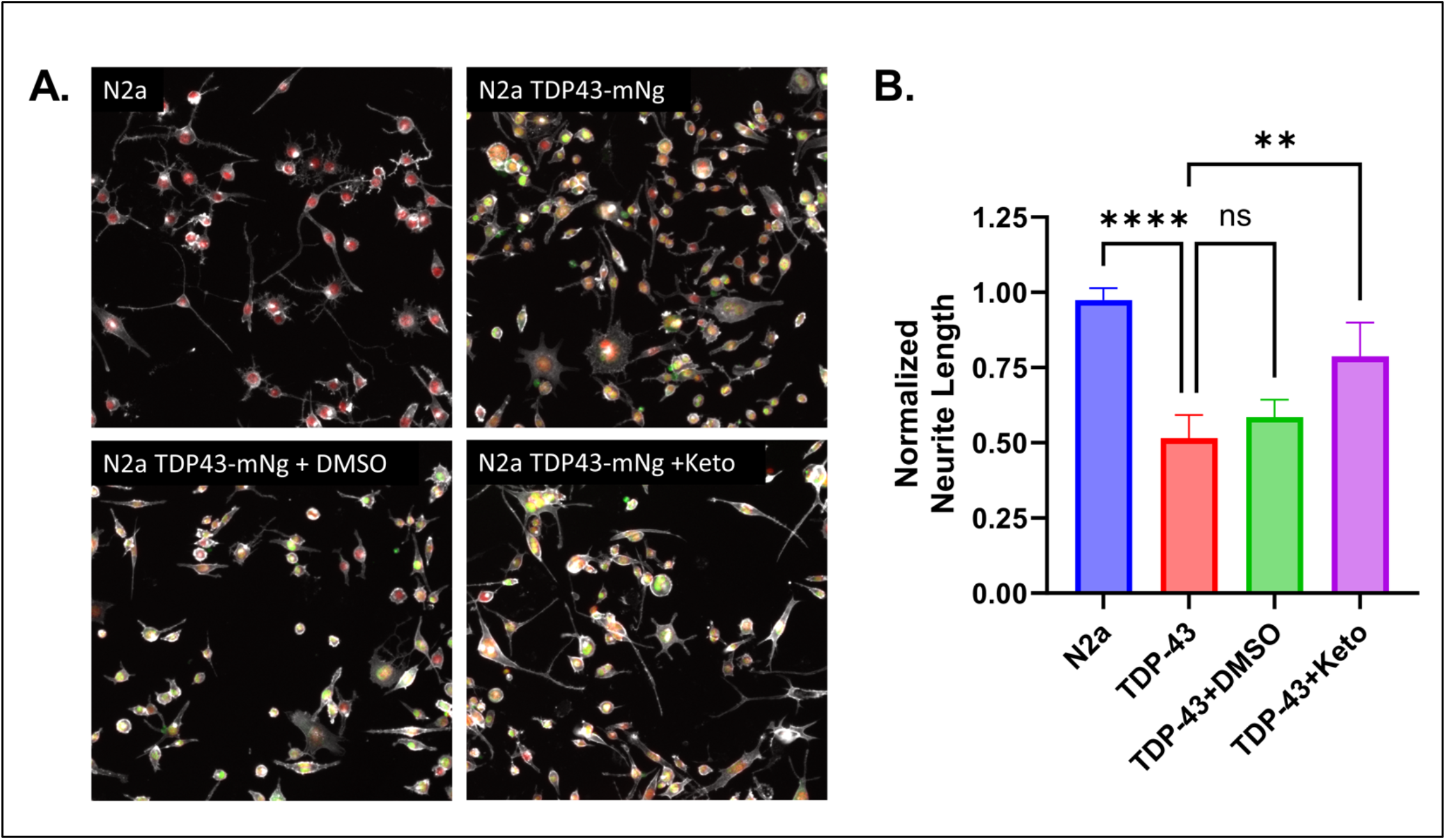
Neurite growth assay in retinoic acid differentiated normal and stable TDP-43-mNg expressing N2a cells. **(A)** Fixed cell fluorescence imaging of N2a and N2a TDP-43-mNg (untreated, DMSO and 10 μM ketoconazole) stained with phalloidin rhodamine (gray pseudo-color), Hoechst (red pseudo-color). mNeonGreen channel shown in green. **(B)** Normalized neurite length (total neurite length normalized to number of cells in each image) for each condition. Statistics shown are two-way ANOVA multiple comparisons to untreated TDP-43-mNg cells adjusted with Bonferroni correction. Data shown are mean ± SEM from N=3 independent experiments with at least 6 images per condition for each run.

### Ketoconazole rescues sorbitol-induced puncta formation and cytoplasmic mislocalization of TDP-43

With ketoconazole demonstrating a positive response in a TDP-43 proteinopathy phenotype assay, we next explored whether it could prevent the formation of TDP-43 puncta and mislocalization upon cellular stress. Sorbitol induced hyperosmotic stress has been widely used to elicit TDP-43 stress response, and recently it was shown to disrupt TDP-43 NTD-interactions, leading to puncta formation and cytoplasmic mislocalization^14,35,36^. Most sorbitol induced hyperosmotic stress assays have used very high levels of insult (e.g., 400 mM), but recent findings suggest lower concentrations are more effective at inducing mislocalization^37^. We first optimized the sorbitol treatment conditions to establish the minimal perturbation that would result in significant changes in FRET for both FL and ΔNTD biosensor while also inducing the formation of TDP-43 puncta and cytoplasmic mislocalization. Supplemental Figure 9A-B shows that 0.1 M sorbitol is sufficient to induce puncta formation and mislocalization of TDP-43; and results in a similar FRET increase relative to vehicle for both FL and ΔNTD biosensors. This consistent FRET change suggests that sorbitol increases NTD-independent TDP-43 assemblies, providing an ideal paradigm to test whether stabilizing NTD-dependent conformations can block TDP-43 cytoplasmic mislocalization and aggregation.

Micrograph images in Figure 5A show that after 20 hours of treatment, sorbitol causes widespread TDP-43 puncta and mislocalization, which is mitigated by 10 *μ*M ketoconazole. Quantification of fluorescence imaging shows a statistically significant increase TDP-43 cytoplasmic mislocalization relative to DMSO-only control, which is significantly reduced by ketoconazole (Figure 5B). In addition, Figure 5C shows that ketoconazole is also able to rescue the doubling in sorbitol-induced TDP-43 puncta at 20 hours to DMSO treatment levels. Supplemental Figure 10 show the treatment design, time traces grouped by vehicle/sorbitol or by DMSO/ketoconazole treatments, as well as a bar plots summarizing all conditions at the final timepoint.

**Figure 5.**
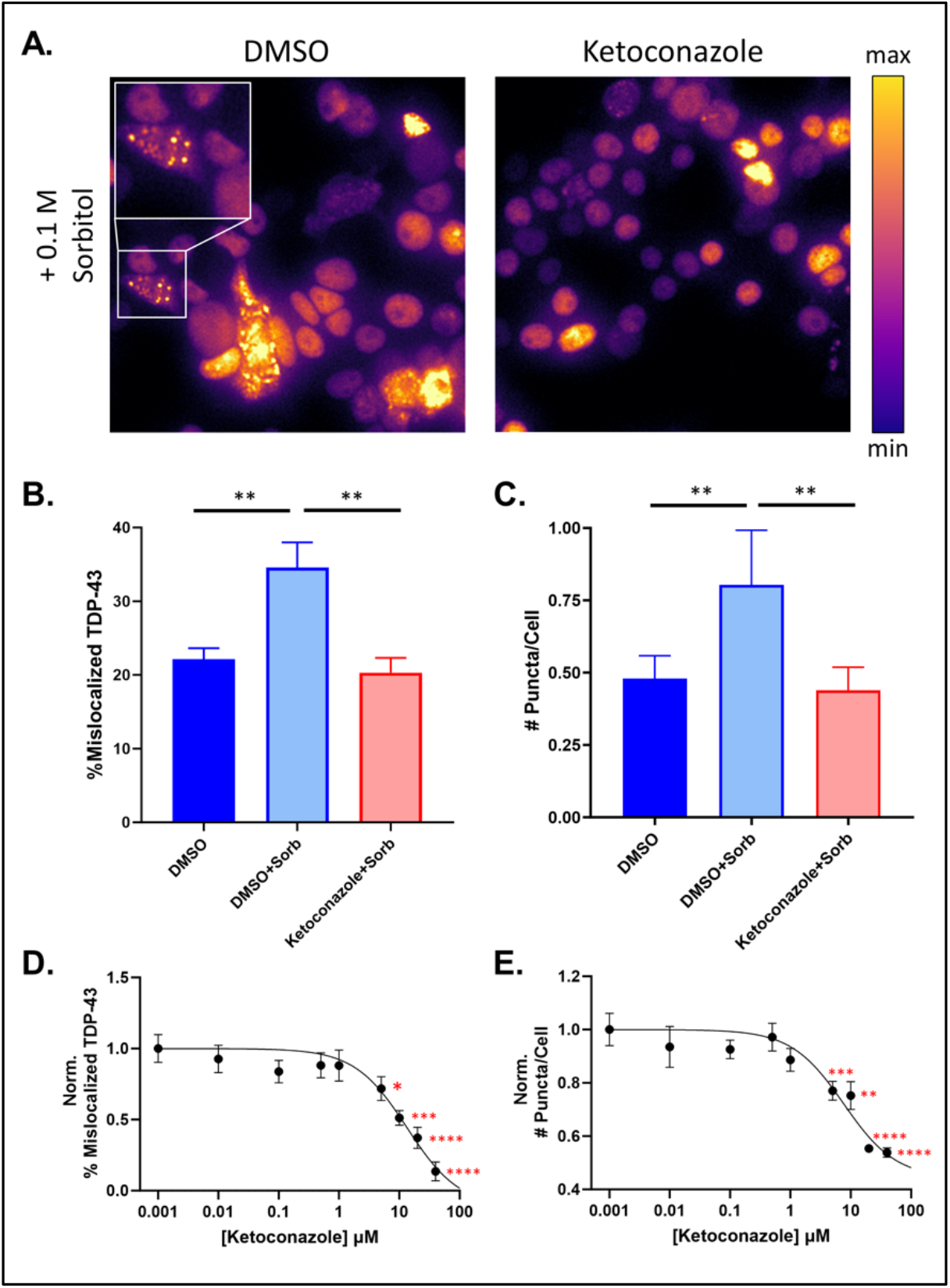
Ketoconazole rescues sorbitol-induced TDP-43 puncta formation and cytoplasmic mislocalization. **(A)** Fluorescence live cell imaging of HEK293T cells expressing FL TDP-43-mNg treated with 0.1 M sorbitol under DMSO or 10 *μ*M ketoconazole treatments. Green fluorescence was mapped to a pseudo-color LUT. **(B)** Endpoint quantification of puncta per cell. **(C)** Endpoint quantification of TDP-43 cytoplasmic mislocalization. Statistics shown in (B-C) are two-way ANOVA multiple comparisons to DMSO-only control (solid blue) with Bonferroni correction (*p < 0.05, **p < 0.01). Supplemental Figures 8-10 show the full dataset of puncta and mislocalization experiments (time traces and drug-only effects). **(D)** Normalized mislocalized TDP-43 as a function of ketoconazole concentration (IC50 = 13.42 *μ*M). **(E)** Normalized number of puncta per cell as a function of ketoconazole concentration (IC50 = 7.65 *μ*M). Statistics shown in (D-E) are ordinary one-way ANOVA multiple comparisons to DMSO-only controls with Bonferroni correction (*p < 0.05, **p < 0.01, *** p < 0.001, ****p < 0.0001). Data shown are mean ± SEM from N=3 independent experiments.

Ketoconazole had a strong effect in both puncta and mislocalization assays; and thus, we tested whether dosing in the nanomolar to micromolar range had a titratable effect on these two phenotypes. Figure 5D-E shows that ketoconazole has a dose responsive effect on mislocalization (IC_50_=13.42μM) and puncta formation (IC_50_=7.65μM) under sorbitol treatment and is able to rescue mislocalization to a greater extent than puncta formation.

### Ketoconazole rescues TDP-43 induced SREPB2 deficiency

Next, we explored the potential MOA of ketoconazole. Ketoconazole is a known binder and inhibitor of the P450 cytochrome lanosterol 14-alpha demethylase (CYP51A), which converts lanosterol to cholesterol in the last enzymatic step of the cholesterol biosynthesis pathway^38^. Inhibition of CYP51A via ketoconazole has been shown to upregulate the master regulator of the cholesterol biosynthesis pathway SREBP2 as a compensatory mechanism to restore homeostatic cholesterol levels^38,39^.

Recently, SREBP2 and the cholesterol biosynthesis pathway have been implicated in the pathogenesis of ALS and FTD^40-42^. SREBP2 acts as a transcription factor when cleaved and regulates the expression of multiple enzymes in the cholesterol biosynthesis pathway via binding to sterol-regulated elements in DNA promoter regions, where inhibition of its activity can have detrimental effects on overall sterol homeostasis and myelination^40,41^. This effect has been shown in HEK293T cells, spinal cord tissue of A315T mutant TDP-43 mice, mouse oligodendrocytes, and cerebrospinal fluid (CSF) samples of ALS patients^40,41^.

Using quantitative reverse transcription polymerase chain reaction (RT-qPCR) we examined whether our biosensor model recapitulates pathological SREBP2 downregulation and if ketoconazole is able rescue this deficit. TDP-43 overexpression caused a significant 2-fold reduction in SREBP2 level relative to untransfected HEK293T cells (Figure 6A; see Supplemental Figure 11 for confirmation of increased TDP-43 protein expression). Treatment with ketoconazole was able to partially rescue this phenotype by increasing SREBP2 levels 1-fold higher relative to untreated TDP-43 overexpressing cells. Overexpression of unlabeled TDP-43 did induce a downregulation of endogenous TDP-43 (detected with 3’-UTR specific primers that do not detect mRNA transcribed from the TDP-43 plasmid), which is expected and consistent with TDP-43’s known autoregulatory mechanism (Figure 6B)^43^, with ketoconazole not showing an additional effect on TDP-43 levels.

**Figure 6.**
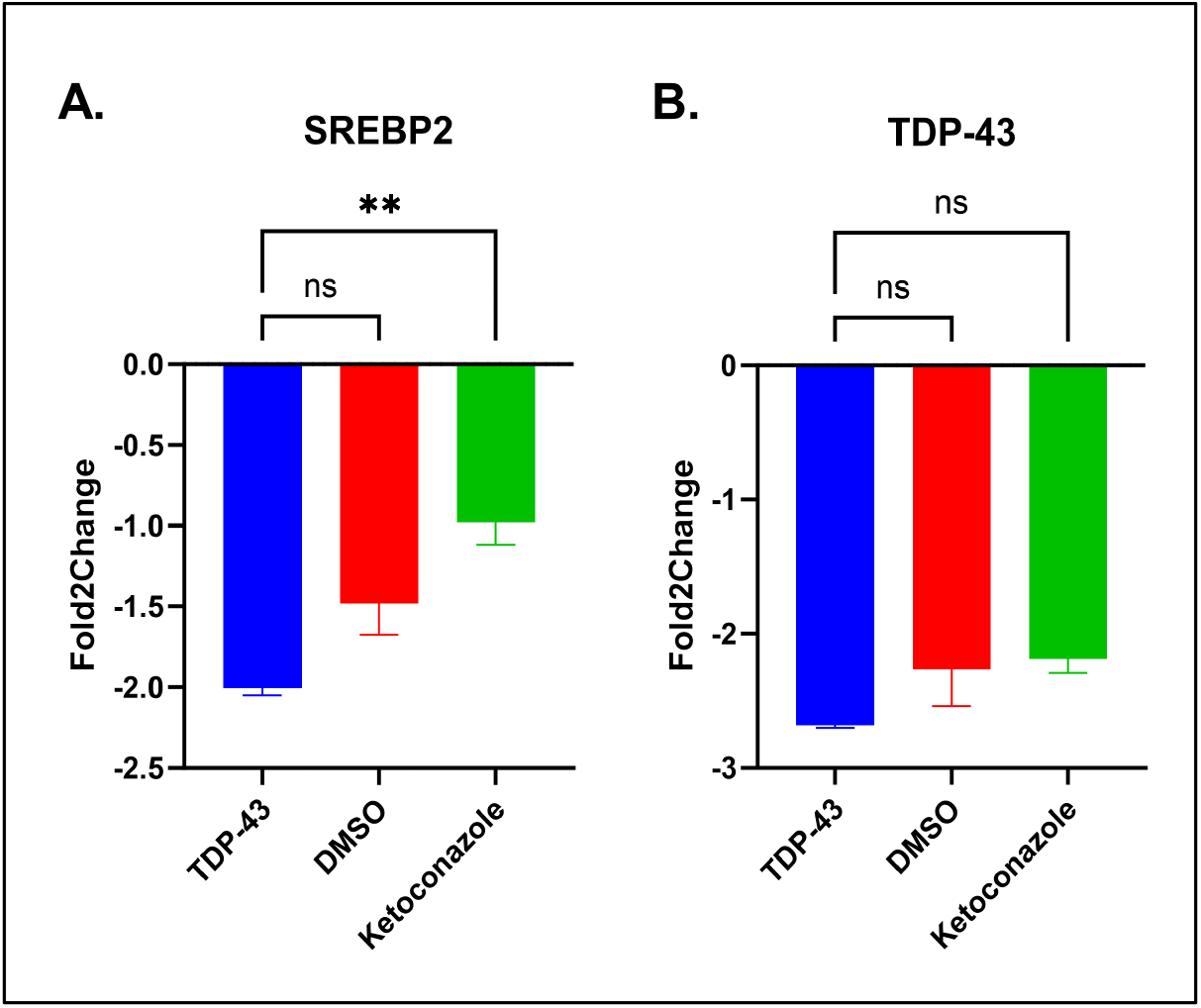
RT-qPCR assay probing for endogenous SREBP2 and TDP-43 under TDP-43 overexpression. **(A)** Fold2Change of endogenous SREBP2 mRNA in untreated, DMSO-treated or ketoconazole-treated HEK293T cells. **(B)** Fold2Change of endogenous TDP-43 mRNA in untreated, DMSO-treated or ketoconazole-treated HEK293T cells. Fold2Changes were calculated relative to non-transfected HEK293T cells and used GAPDH as a housekeeping gene. Statistics shown are one-way ANOVA multiple comparisons with Bonferroni correction against untreated TDP-43 overexpressing cells (blue bar, **p < 0.01). Data shown are mean ± SEM from N=3 independent experiments.

To further explore this potential MOA, we mined De Abrew et. al.’s publicly available transcriptomic data of four cancer cell lines (HepG2, MCF7, HepaRg and Ishikawa cells) treated with 34 different small molecules, including ketoconazole^44^. Indeed, at 10 *μ*M dosing (matching our FLT-FRET experimental conditions) we found that ketoconazole increased the expression of SREBP2, along with 14 out of 15 of its downstream regulated mRNAs coding for proteins involved in the isoprenoid and cholesterol biosynthesis pathway. Interestingly, TDP-43 is known to bind to and regulate 11 of these 15 proteins at the mRNA level (Supplemental Figure 12A-B)^41^. As a negative control, a similar analysis of amoxicillin, which was not a hit in our FLT-FRET screens, does not cause any significant change relative to vehicle on the levels of these mRNAs (Supplemental Figure 12A-B).

### Ketoconazole improves motor behavior in TDP-43 overexpressing C. elegans

We then explored ketoconazole’s ability to rescue a TDP-43 induced pathological phenotype in a C. elegans model of TDP-43 proteinopathy. The OW1601 strain developed by Koopman et. al. expresses human TDP-43 pan-neuronally and exhibits motor deficits that are accompanied by formation of TDP-43 insoluble aggregates^45,46^. Both control (OW1603) and hTDP-43 (OW1601) animals were grown in DMSO or ketoconazole (10 or 50 μM) from synchronized L1 to L4 stage. Motor deficits were monitored via a swimming assay which tracked body bends per 30 seconds (BB per 30s).

Figure 7 shows the hTDP-43 strain has a significantly reduced number of BB per 30s while swimming relative to controls. Ketoconazole treatment improves motor function in the hTDP-43 animals more than two-fold relative to DMSO (∼16% to ∼40% of healthy controls). Importantly, ketoconazole does not enhance the motor function of control animals, suggesting a TDP-43-specific effect in the hTDP-43 strain.

**Figure 7.**
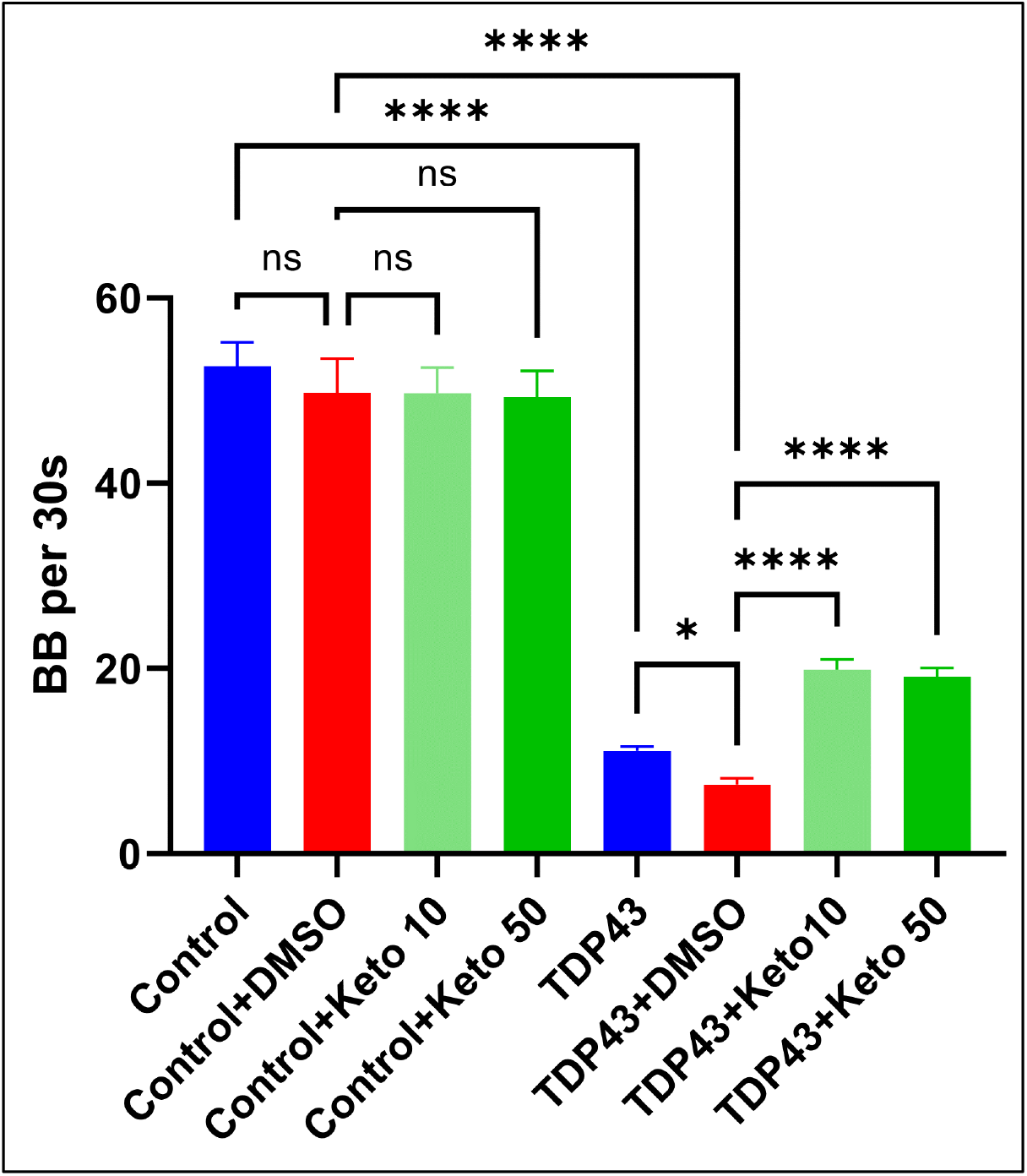
Ketoconazole improves motor dysfunction in human TDP-43 overexpressing C. elegans OW1601 strain. **(A)** BB per 30s for L4 stage control OW1603 strain and hTDP-43 strain OW1601 under DMSO or 10 μM ketoconazole treatment. Statistics shown are one-way ANOVA multiple comparisons with Bonferroni correction (*p < 0.05, ****p < 0.0001). Data shown are mean ± SEM from N=3 batches containing at the minimum 10 animals.

## Discussion

Our screening platform demonstrates the ability to target TDP-43 NTD-dependent interactions in living cells with a small molecule, validating the therapeutic premise presented by Oiwa et. al^17^. Our biosensors are amenable to HTS and offer a 16-fold improvement in precision over fluorescence intensity methods. Our counter-screen strategy using FL and ΔNTD biosensors identified the hit compound ketoconazole, which rescued TDP-43 dependent deficits in neurite outgrowth of differentiated neurons and motor behavior in C. elegans, sorbitol-induced TDP-43 puncta formation and cytoplasmic mislocalization, and downregulation of the master cholesterol regulator SREBP2.

Ketoconazole is a known binder and inhibitor of CYP51A, an enzyme in the cholesterol synthesis pathway. Interestingly, ropinirole, an ALS drug currently in clinical trials, has also been shown to improve ALS patient derived iPSC motor neurons phenotypes via modulation of the cholesterol biosynthesis pathway^6^, yet the precise link to TDP-43 is not yet known. Our platform’s ability to identify ketoconazole as a modulator of similar disease phenotypes, potentially via an analogous MOA, suggests a benefit to monitoring and targeting TDP-43 NTD interactions.

Although ketoconazole was the only compound with no significant ΔNTD biosensor response — passing the stringent ΔNTD counter-screen filter — some compounds displayed a reduced ΔNTD biosensor response but under the conditions tested were not completely ΔNTD independent. In total, 14 of the 23 FL TDP-43 hits had a reduced significance in the ΔNTD counter-screen (Supplemental Figure 7). Two such hits, ginsenoside Rb1 and rifabutin, also have a known MOA related to the cholesterol biosynthesis pathway^47,48^. Both ginsenoside Rb1 and rifabutin had a partial NTD-dependent ΔFLT effect (Supplemental Figure 13) and were able to partially rescue TDP-43 mislocalization, puncta formation due to sorbitol (Supplemental Figure 14) and SREBP2 levels due to TDP-43 overexpression (Supplemental Figure 15). Another partial NTD-dependent compound, radotinib, is a C-abl kinase inhibitor — a MOA that is also being pursued for ALS^49,50^ — is currently being explored as a potential Parkinson’s disease therapeutic^49^. These results suggest that broadening our counter-screen criterion to include partial NTD-dependent molecules may enrich for additional compounds with activity towards TDP-43.

Part of the hit criterion that was used for this study was to focus on compounds that reduced NTD-dependent FLT (increased FRET and therefore increased NTD-dependent interactions). Six hit compounds displayed an increased in FLT (reduced FRET) with four such compounds having an NTD-dependent response (Supplemental Figure 7C). Indeed, the hit auranofin was previously identified in a screen targeting TDP-43 aggregation, suggesting that compounds like these may warrant further investigation as possible tool compounds that alter the distribution of toxic TDP-43 assemblies.

Our domain specific counter-screen strategy could be expanded to investigate the other domains of TDP-43. Modifications to the RRMs or CTD, as well as the nuclear localization and export sequences, could be introduced to bias the ensemble of TDP-43 assemblies in the biosensor cells to specific states or interactions along TDP-43’s aggregation cascade. For example, by screening against FL and a ΔCTD TDP-43 under proteostatic stress conditions, our approach could identify compounds that reduce FRET (reducing CTD-CTD interactions) and target the CTD specifically. In addition, this platform can be utilized to understand the effects of ALS-related mutations (in TDP-43 and other proteins such as FUS and C9orf72) on the NTD-mediated oligomerization of TDP-43. Similarly, this approach can be used to explore whether ALS-risk factors such as neuroinflammation, environmental toxins or commonly prescribed drugs can detrimentally modulate TDP-43 assemblies. For example, our platform identified the erdafitinib, an FGFR1 inhibitor commonly prescribed for bladder cancer^51^, as an NTD-independent strong inducer of TDP-43 pathology. Overall, we speculate that our platform can be used to determine whether loss of NTD interactions is a converging node for divergent ALS-triggering factors.

## Methods

### Cell culture

HEK293T cells (ATCC) were maintained in full Gibco DMEM media (10% FBS, 1% Pen/Strep, 1% GlutaMax) at 37 °C, 5% CO2, and humidity control. All transfections were carried out using Thermo Fisher Lipofectamine 3000 at a DNA:P3000:L3000 ratio of 1:2:3 (*μ*g DNA: *μ*L P3000: *μ*L L3000), with 2.5 *μ*g DNA per 1e6 seeded cells.

N2a cells (ATCC) were maintained in full EMEM media (10% FBS, 1% Pen/Strep, 1% GlutaMax, 1% NEAAs) and differentiated in 0.5% FBS with 10 *μ*M retinoic acid. Stable WT TDP-43-mNg expressing N2a cells were generated by transfection with Lipofectamine 3000, followed by selection with Zeocin and cell sorting using flow cytometry.

### Western blotting

Cells were harvested using trypsin (Gibco) and washed in PBS (Sigma-Aldrich) once prior to lysis with (ultrapure water, TNE buffer, 1% SDS, 0.5% DOC, 0.5% NP-40, phosphatase and kinase inhibitors). Samples were incubated on ice for 10 minutes, sonicated at 50% amplitude for 3x 3s pulses, boiled at 95 °C for 10 minutes and spun down at 21,300g for 10 minutes. Using the supernatant, bicinchoninic acid assays (BCA, Thermo Scientific) were conducted to quantify total protein content and samples prepared in 1x Laemmli buffer (BioRad) with beta-mercaptoethanol.

SDS-PAGE gels were run at 120V using either 12% or 4-20% BioRad gels with 10-20 *μ*g of protein loaded in each well. Gels were transferred into nitrocellulose membranes using the BioRad TurboTransfer protocol and stained for total protein using Ponceau S. Finally, blots were probed with TDP-43 (Proteintech 10782-2-AP), mNeonGreen (Cell Signaling Technology E6M3D), mCherry (Cell Signaling Technology E5D8F) primary antibodies at 1:2000 dilution overnight at 4 °C and detected using HRP-conjugated secondary antibodies at 1:10000 dilution imaged with SuperSignal West Pico PLUS chemiluminescent substrate (Thermo Scientific).

### Intact cell cross-linking

DSG cross-linking was conducted by incubating 1e6 PBS-washed cells resuspended in 500 *μ*L with 200 *μ*M DSG (Sigma Aldrich) for 30 minutes under gentle rotation. Next, the cross-linking reaction was quenched with 20 mM Tris (pH 7) for 15 minutes at room temperature. After quenching, cells were spun down and processed for western blotting as described above.

### Cell preparation and FLT-FRET measurements

For all FLT-FRET experiments, cells were transfected with 0.5 *μ*g/1e6 cells TDP-43-mNg (donor-only), 0.5 *μ*g/1e6 cells TDP-43-mNg + 0.5 *μ*g/1e6 cells TDP-43-mCh (1:1 FL), 0.5 *μ*g/1e6 cells TDP-mNg + 0.5 *μ*g/1e6 cells TDP-43-mCh ΔNTD (1:1 ΔNTD) or 0.5 *μ*g/1e6 cells mCh-Linker-mNg complemented with empty vector plasmid to reach 2.5 *μ*g DNA/1e6 cells. After 24 hours of transfection, cells were lifted using trypsin, washed in PBS twice and resuspended at a final concentration of 1e6 cells/mL. 50 *μ*L of cells were added to black-bottom 384-well plates where FLT measurements were collected.

FLT measurements were made in a fluorescence lifetime plate reader (Fluorescence Innovation, USA) equipped with a 473 nm laser, 488 dichroic, and dual channel detection at 517/20 (primary) and 535/6 (secondary). ^30^. FLT for each well is calculated by fitting a single exponential decay function to the time-resolved fluorescence waveforms as described previously^30^. FRET efficiency was then calculated using Equation 1.

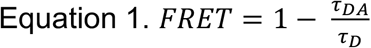

### Cell preparation and FDA-approved Selleck library FLT-FRET screens and dose-responses

Cells were transfected with full-length and linker-only biosensors in 15 cm plates using the same DNA concentrations listed previously. After 24 hours of transfection, cells were lifted, washed in PBS twice, passed through a cell strainer, and resuspended to 1e6 cells/mL. The Selleck library used consists of 2684 FDA-approved small molecules pre-dispensed across three separate 1536-well plates, which include empty and DMSO wells as negative controls. After equilibrating the plates to room temperature, 5 *μ*L of cells were dispensed in each well using a Thermo Fisher Multidrop Combi for a final drug concentration of 10 *μ*M. Plates were incubated at room temperature for 2 hours prior to FLT-FRET measurements. TDP-43 and linker unique hits were determined by finding 3SD hits that appeared in each independent screen, flagging interfering compounds with a spectral similarity filter as described previously^30^. TDP-43 unique hits were defined as hits from the TDP-43 screen that did not show up as hits in the linker screen.

TDP-43 reproducible and unique hits were then tested by conducting FLT-FRET measurements at increasing concentrations (∼156 nM to 10 *μ*M). The exact same steps and incubation times were followed for dose responses, with the only difference being that these experiments were conducted in 384-well plates pre-dispensed with drugs (see chemical vendors in Supplemental Table 1) using a Perkin Elmer Flexdrop liquid dispenser.

### Neurite length assay

Neuro2a (N2a, ATCC) cells were plated at a density of 70K cells/well in a 24-well plate. After 6 hours, 10 *μ*M retinoic acid differentiation media was added to the wells. After 24 hours of incubation, 10 *μ*M ketoconazole or DMSO was added for an additional 24 hours. Wells were washed in PBS, fixed in 4% paraformaldehyde, permeabilized with 0.5% Triton-X and stained with Hoechst and phalloidin rhodamine (Cytoskeleton, Inc) following the manufacturers protocol.

### TDP-43 puncta formation and mislocalization imaging and analysis

HEK293T cells were seeded at 0.16e6 cells per well in 24-well plates. After 24 hours of incubation, each well was transfected with 0.08 *μ*g TDP-43-Ng (supplemented with 0.32 *μ*g empty vector for a total of 0.4 ug DNA per well, 2.5 *μ*g DNA/1e6 cells). After 24 hours of incubation, cells were pre-treated with 0.2% DMSO control and 10 *μ*M ketoconazole, ginsenoside Rb1 and rifabutin for 2 hours. After 1.5 hours, Hoechst stain was added at a 1:50k dilution and incubated for 30 minutes. Next, ultra-pure water control or 0.1 M sorbitol were added to wells. Plates were imaged for nuclear, TDP-43-mNg and brightfield channels continuously under environmental control (37 °C, 5% CO2, 15% O2 and humidity control) every hour for 20 hours since addition of drug in a Molecular Devices Pico ImageXpress Pico microscope.

Images were analyzed using MetaXpress High Content Image Analysis software with two custom analysis workflows that quantify number of puncta per cell and cytoplasmic mislocalization based on intensity thresholding and nuclear masking using the Hoechst channel.

### Mining of ketoconazole treatment transcriptomics data

Differences in expression of SREBP2 and its regulated genes between vehicle and ketoconazole or amoxicillin in HepG2, MCF7, HepaRg and Ishikawa cells were calculated using the data deposited by De Abrew et. al. on NCBI’s Gene Expression Omnibus (GEO)^44^. The integrated GEO2R analysis suite was utilized to calculate log2FC expression changes of SREBP2-regulated genes in the cholesterol biosynthesis pathway.

### Real-Time Quantitative PCR of endogenous TDP-43 and SREBP2

HEK293T cells were seeded at 0.8e6 cells per well in 6-well plates and incubated for 24 hours. After incubation, cells were transfected with 2 *μ*g of unlabeled full-length TDP-43 (2.5 *μ*g DNA/1e6 cells). Two hours after transfection, cells were treated with 10 *μ*M ketoconazole, ginsenoside Rb1 and rifabutin (as well as 0.2% matching DMSO control) and incubated for an additional 24 hours.

Total RNA for each condition was isolated using Qiagen RNAeasy extraction protocol. After quality validation via 280/260 measurements, total RNA was reversed transcribed using Thermo Fisher High-Capacity RNA-to-cDNA kit. PCR reactions probing for endogenous GAPDH (housekeeping), TDP-43 and SREBP2 were prepared following the Thermo Fisher SYBR Green qPCR protocol and using the following specific primers (IDT) for endogenous mRNA:

GAPDH

Forward: 5’-AATGGGCAGCCGTTAGGAAA-3’

Reverse: 5’-GCGCCCAATACGACCAAATC-3’ TDP-43:

Forward: 5′-GAGAAAAGGAGAGAGCGCGT-3′

Reverse: 5′-GGGGTAGGGGGAGTACAAGT -3′

SREBP2:

Forward: 5’-TGTGTCCTCACCTTCCTGTGCCT-3’

Reverse: 5’-TCCAGTCAAACCAGCCCCCAGA-3’

RT-qPCR reactions were conducted using a Roche LightCycler 96 plate reader and analyzed using Roche’s PCR analysis software. Expression was quantified using the 2^-ΔΔCt^ method relative to untreated/untransfected HEK293T cells^52^.

### C. elegans motor behavior analysis via swimming assay

Control (OW1603, dvIs15 [unc-54(vector) + mtl-2::GFP]) and human TDP-43 overexpressing (OW1601, dvIs62 [snb-1p::hTDP-43/3’ long UTR + mtl-2p::GFP]) strains were obtained from Ellen A. A. Nollen from the University of Groningen and long-term stored at the *Caenorhabditis* Genomics Center (CGC) at the University of Minnesota. Animals were grown on nematode growth medium (NGM) plates at 21°C. Drug plates were prepared using NGM agar supplemented with 0.2% DMSO, 10 or 50 *μ*M ketoconazole in a 6 cm plate. Solidified plates were seeded with OP50 in 0.2% DMSO, 10 or 50 *μ*M ketoconazole.

Animals were synchronized at L1 stage following established protocols^53^. Hatched L1 larvae were distributed onto NGM agar drug plates (∼10 animals per well in triplicate wells per each condition). Worms were incubated at room temperature for 3 days. Worms were collected, washed in M9 buffer and plated in 12-well NGM agar plates containing sufficient M9 buffer for monitoring swimming behavior. Animals were allowed to equilibrate for 1 minute before recording 30 second videos. The video recordings were processed and analyzed to quantify BB per 30s using the wrMTrck plugin in ImageJ.

### Statistical analysis

Multiple comparisons were conducted via ordinary one-way or two-way ANOVAs with post-hoc Bonferroni corrections for multiple comparisons. Single comparisons were conducted via one sample T-tests or unpaired two-tailed T-tests. All statistical analysis was done in GraphPad Prism 9.

## Supplemental Material

Supplemental data includes table summarizing TDP-43 unique hits, biosensor expression control western blots, FLT dose response curves, crosslinking western blots and quantification, sorbitol-induced puncta and mislocalization time traces, and full RT-qPCR and transcriptomic analysis.

## Data Availability

The datasets generated and/or analyzed during the current study are available from the corresponding author on reasonable request.

## Code Availability

Fluorescence lifetime analysis was conducted using a single-exponential fitting script written in MATLAB. All scripts are available from the corresponding author on reasonable request.

## Acknowledgments

The authors disclose receipt of the following financial support for the research, authorship, and/or publication of this article: The C. elegans experiments were supported by the *Caenorhabditis* Genetics Center (CGC), which is funded by the NIH office of Research Infrastructure Programs (P40 OD10440). C. elegans strains (OW1601 and OW1603) were a generous gift from Ellen A. A. Nollen at the University of Groningen, and have been deposited at the CGC.

## Author Contributions

N.N.K., A.R.B., and J.N.S. conceived of and directed the study; N.N.K. conducted and directed all experimental work; M.M. conducted plasmid DNA preparations and western blotting; N.V. and E.E.L. assisted with western blotting and fluorescence imaging; L.C. assisted with C. elegans experimental design and edited the manuscript, N.N.K., A.R.B., and J.N.S wrote and edited the manuscript.

