## Supplemental_Information for "Fluorescence lifetime-based FRET biosensors for monitoring N-terminal domain interactions of TDP-43 in living cells: A novel resource for ALS and FTD drug discovery"

Noah Nathan Kochen[1*], Marguerite Murray[1], Nagamani Vunnam[1], Elly E. Liao[1], Lihsia Chen[2], Anthony R. Braun[1‡,] and Jonathan N. Sachs[1‡]
[1]Department of Biomedical Engineering, University of Minnesota, Minneapolis, MN 55455
[2]Department of Genetics, Cell Biology and Development, University of Minnesota, Minneapolis, MN 55455

[*]First author; ‡ co-corresponding author

**Keywords**
TDP-43
Neurodegeneration
FRET

### Supplementary Information

Table of Contents:

◊ Supplemental Table 1. Summary of FDA-approved Selleck library TDP-43 unique hits.

◊ Supplemental Figure 1. Comparison of fluorescence intensity and lifetime sensitivity.
◊ Supplemental Figure 2. FRET of FL and ∆NTD at increasing TDP-43 expression and donor:acceptor ratios.
◊ Supplemental Figure 3. Western blot of expression of mNeonGreen and mCherry full-length and ∆NTD TDP-43 biosensors in HEK293T cells.
◊ Supplemental Figure 4. DSG cross-linking summary of HEK293T expressing FL and ∆NTD biosensors.
◊ Supplemental Figure 5. Western blots of un-crosslinked and crosslinked HEK293T cells expressing FL and ∆NTD mNeonGreen constructs.
◊ Supplemental Figure 6. Diagrams of FRET biosensors used in FDA-approved Selleck library screen.
◊ Supplemental Figure 7. ∆FLT responses of all 23 TDP-43 unique hit compounds in FL, ∆NTD and linker biosensors.
◊ Supplemental Figure 8. Live-cell imaging and ∆FLT profile of ketoconazole and erdafitinib.
◊ Supplemental Figure 9. Sorbitol effect on TDP-43 subcellular localization and FRET.
◊ Supplemental Figure 10. Sorbitol-induced TDP-43 puncta formation and mislocalization experimental design and traces for control and ketoconazole treatments.
◊ Supplemental Figure 11. Western blot confirmation of unlabeled TDP-43 overexpression in HEK293T cells for RT-qPCR experiment.
◊ Supplemental Figure 12. Transcriptomics mining of 10 µM ketoconazole treatment on 4 different immortalized cell lines.
◊ Supplemental Figure 13. Live-cell imaging and ∆FLT profile of ginsenoside Rb1 and rifabutin.
◊ Supplemental Figure 14. Sorbitol-induced TDP-43 puncta formation and mislocalization under ginsenoside Rb1 and rifabutin treatment.
◊ Supplemental Figure 15. Full RT-qPCR assay for endogenous SREBP2 and TDP-43 in the presence of DMSO or TDP-43 hits.

| | Drug Name | Delta Lifetime Z-Score | Standard Error | Selleck-Listed Target | Vendor |
|---|---|---|---|---|---|
| 1 | Danthron | -11.621 | 3.951 | N/A | Sigma-Aldrich |
| 2 | Rifapentine | -6.702 | 0.192 | DNA/RNA synthesis | Selleck |
| 3 | Radotinib | -5.867 | 1.165 | BCR-Abl | Selleck |
| 4 | Sennoside A | -5.281 | 0.577 | MAO | Selleck |
| 5 | Erdafitinib (JNJ-42756493) | -5.265 | 0.648 | FGFR | Cayman Chemical |
| 6 | Rifamycin sodium salt | -5.004 | 0.991 | Others | USP |
| 7 | Proanthocyanidins | -4.947 | 0.708 | Others | Selleck |
| 8 | Dithranol | -4.412 | 1.605 | Others | Oakwood Chemical |
| 9 | Levothyroxine sodium | -4.222 | 1.186 | TR-alpha/beta | Sigma-Aldrich |
| 10 | Evans Blue | -3.480 | 1.029 | GluR | Sigma-Aldrich |
| 11 | Nystatin (Fungicidin) | -3.416 | 0.648 | Anti-infection | Cayman Chemical |
| 12 | Diacerein | -3.408 | 1.417 | IL eceptor | Cayman Chemical |
| 13 | Ketoconazole | -3.308 | 0.534 | P450 | CHEM-IMPEX |
| 14 | Rifabutin | -3.120 | 0.062 | Anti-infection | CHEM-IMPEX |
| 15 | Oxytetracycline Dihydrate | -3.024 | 0.234 | Anti-infection | Selleck |
| 16 | Ruboxistaurin (LY333531 HCl) | -2.905 | 1.949 | PKC | Chem Cruz |
| 17 | Ginsenoside Rb1 | -2.747 | 0.275 | Others | Selleck |
| 18 | Auranofin | 3.721 | 2.003 | Others | Selleck |
| 19 | Clindamycin palmitate HCl | 3.856 | 0.619 | Others | Selleck |
| 20 | Trapidil | 4.298 | 0.557 | PDGFR | Selleck |
| 21 | Omeprazole Sodium | 4.450 | 0.215 | Proton pump | Selleck |
| 22 | Allopregnanolone | 6.526 | 3.088 | GABA receptor | Spectrum Chemical MFG |
| 23 | Pomalidomide | 8.543 | 2.405 | TNFalpha | Cayman Chemical |

**Supplemental Table 1. Summary of FDA-approved Selleck library TDP-43 unique hits.** For each hit, the ∆FLT Z-score, standard error, Selleck-reported target and vendor for follow-up experiments are shown.

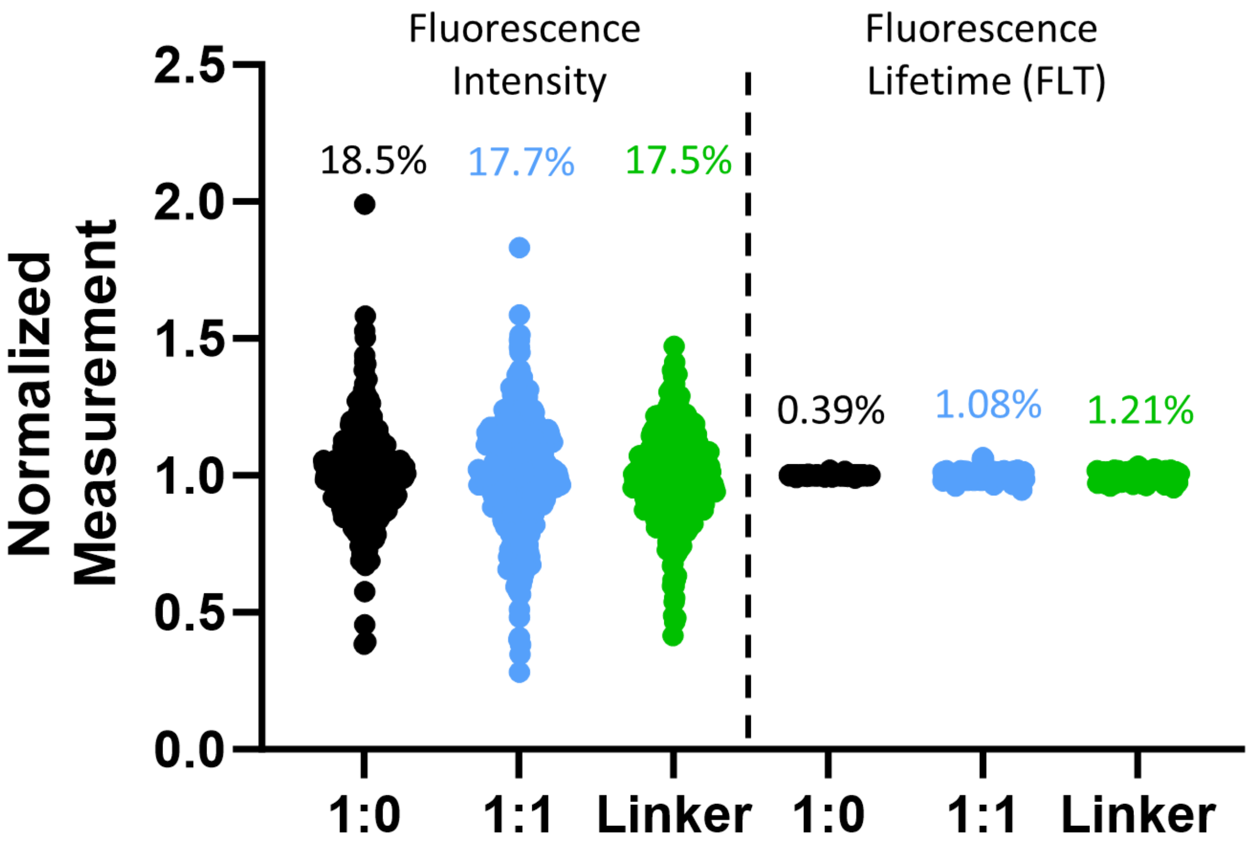


**Supplemental Figure 1. Fluorescence lifetime is 16-fold more sensitive than fluorescence intensity.** Normalized fluorescence intensity and fluorescence lifetime for 473nm excitation channel of TDP-43-mNeonGreen (1:0), TDP-43-mNeonGreen + TDP-43-mCherry (1:1) and mNeonGreen-Linker-mCherry (Linker) constructs used for monitoring TDP-43 FLT-FRET in live HEK293T cells. Percentages indicate % coefficient of variation (%CV) of 384 individual wells of a 1536-well plate.

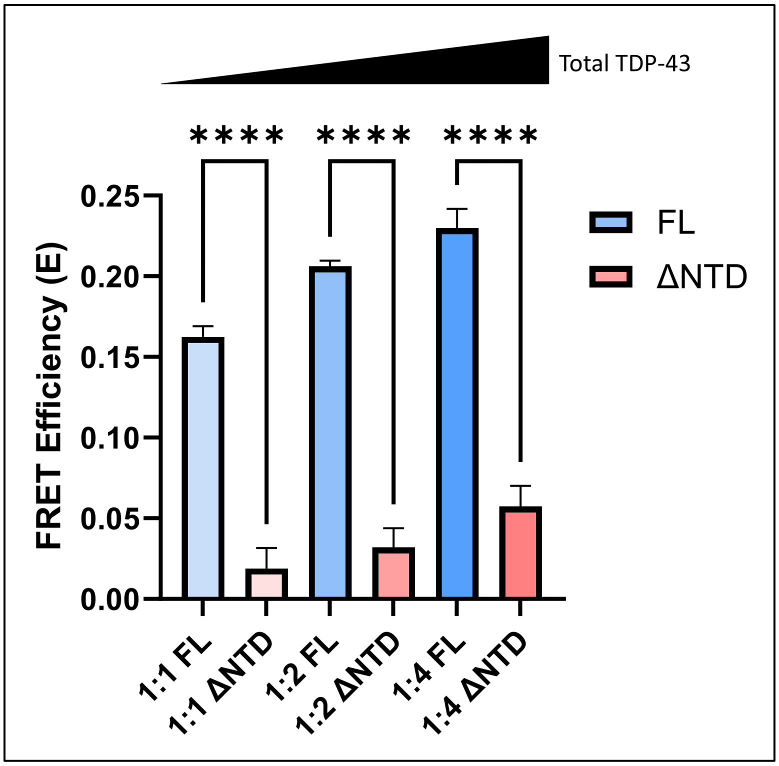


**Supplemental Figure 2. Non-NTD dependent FRET increases with higher levels of TDP-43 biosensor expression.** FRET efficiencies of FL and ΔNTD TDP-43 biosensors at different donor:acceptor ratios (1:1, 1:2, 1:4) and increasing total mass amount of biosensor plasmid transfected (0.8, 1.2, 1.6 μg). Statistics shown are one-way ANOVA multiple comparisons with Bonferroni correction (**** $p < 0.0001$). Data shown are mean ± SEM from N=3 independent experiments.

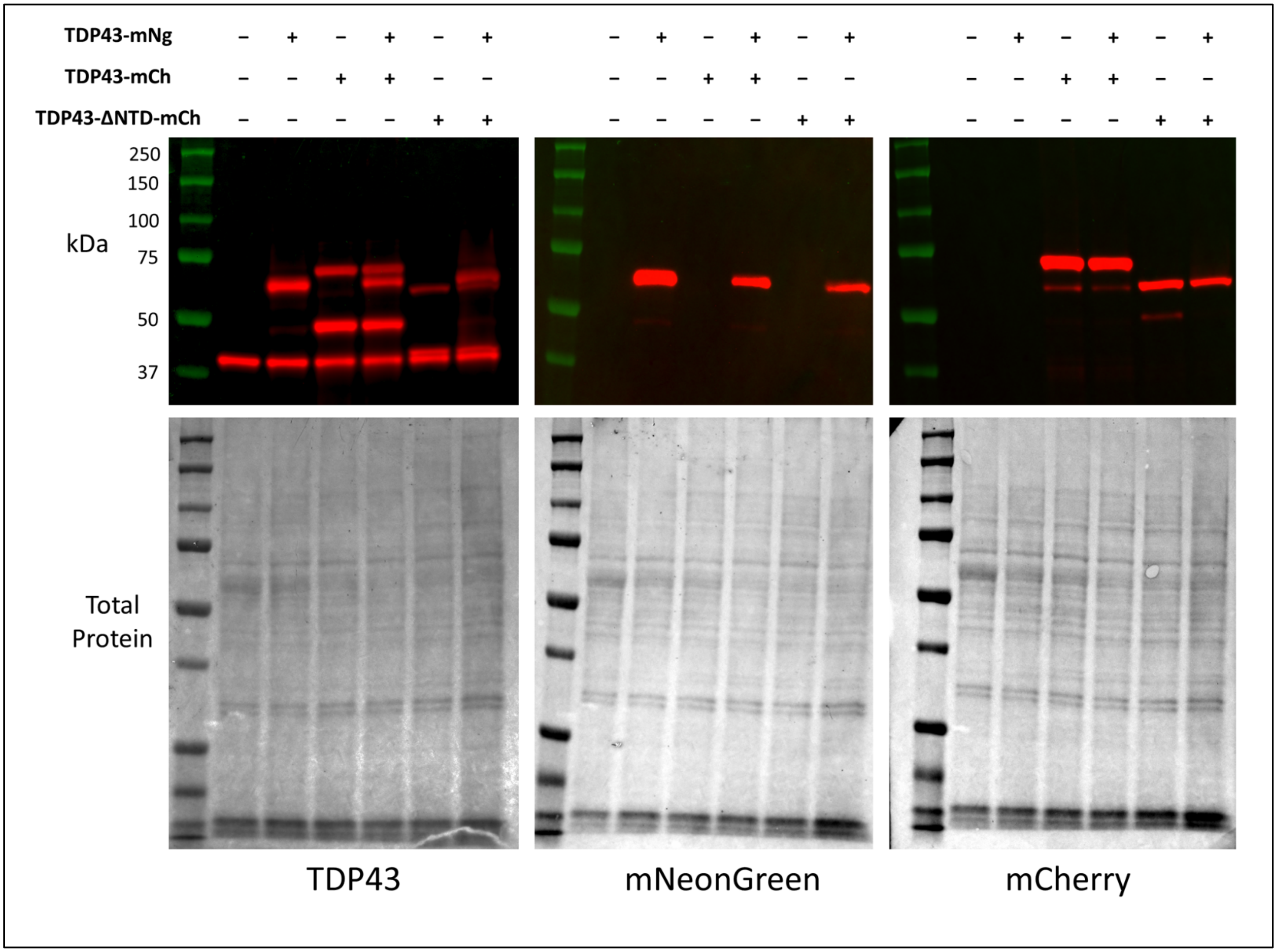


**Supplemental Figure 3. Expression of mNeonGreen and mCherry FL and ΔNTD TDP-43 biosensors in HEK293T cells.** Biosensor expression monitored via western blots probing for TDP-43, mNeonGreen and mCherry.

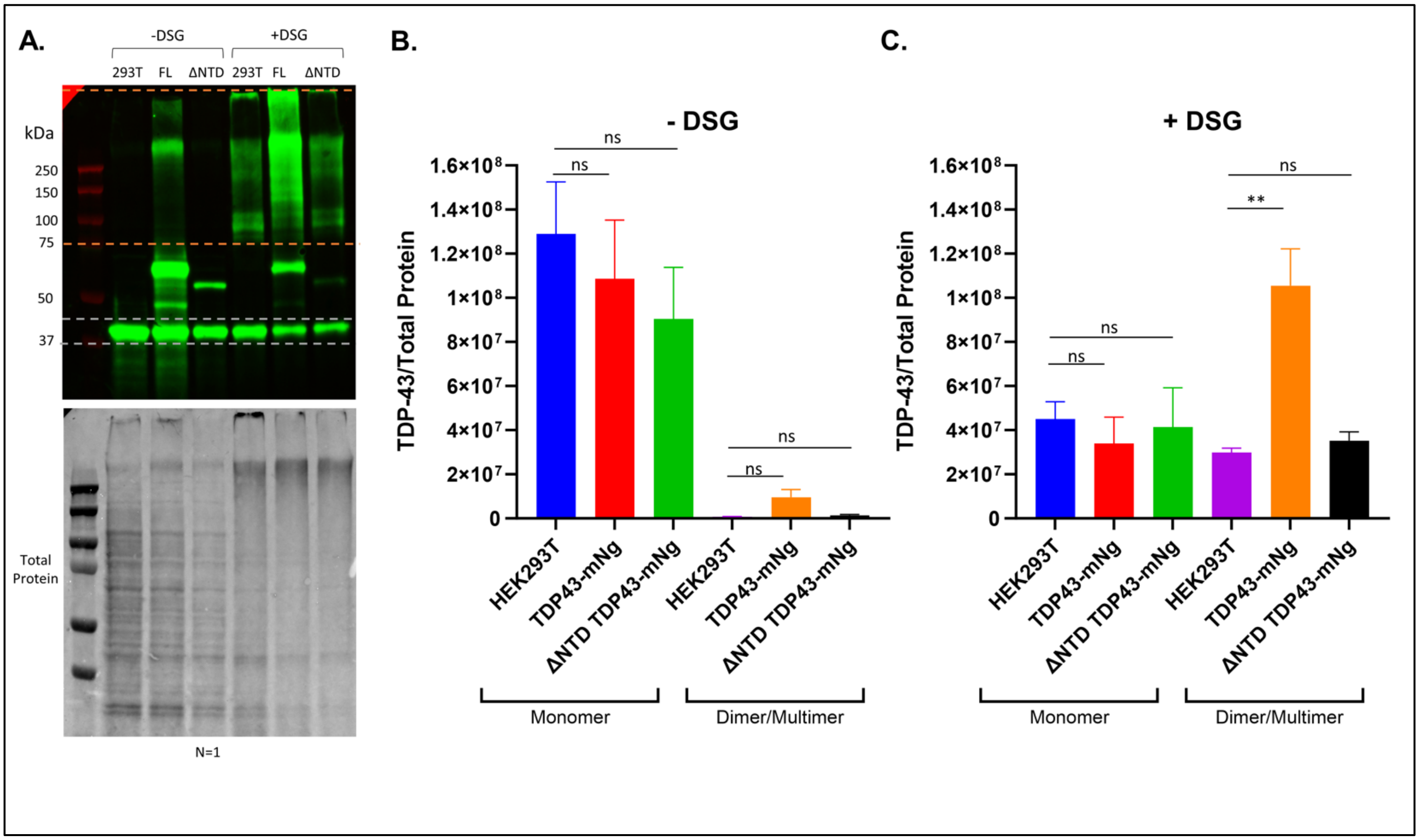


**Supplemental Figure 4. Full-length biosensor expression correlates with higher TDP-43 dimer/multimer levels. (A)** Western blot of HEK293T cells expressing full length and ΔNTD TDP-43 donor biosensor with and without DSG crosslinking. Total protein was stained using Ponceau S. **(B)** Quantification of TDP-43 monomer and dimer/multimer levels in un-crosslinked samples. **(C)** Quantification of TDP-43 monomer and dimer/multimer levels in crosslinked samples. The two additional western blots used for quantification in (B-C) are shown in Supplemental Figure 5. Statistics shown are one-way ANOVA multiple comparisons with Bonferroni correction (**$p < 0.01$). Data shown are mean ± SEM from N=3 independent experiments.

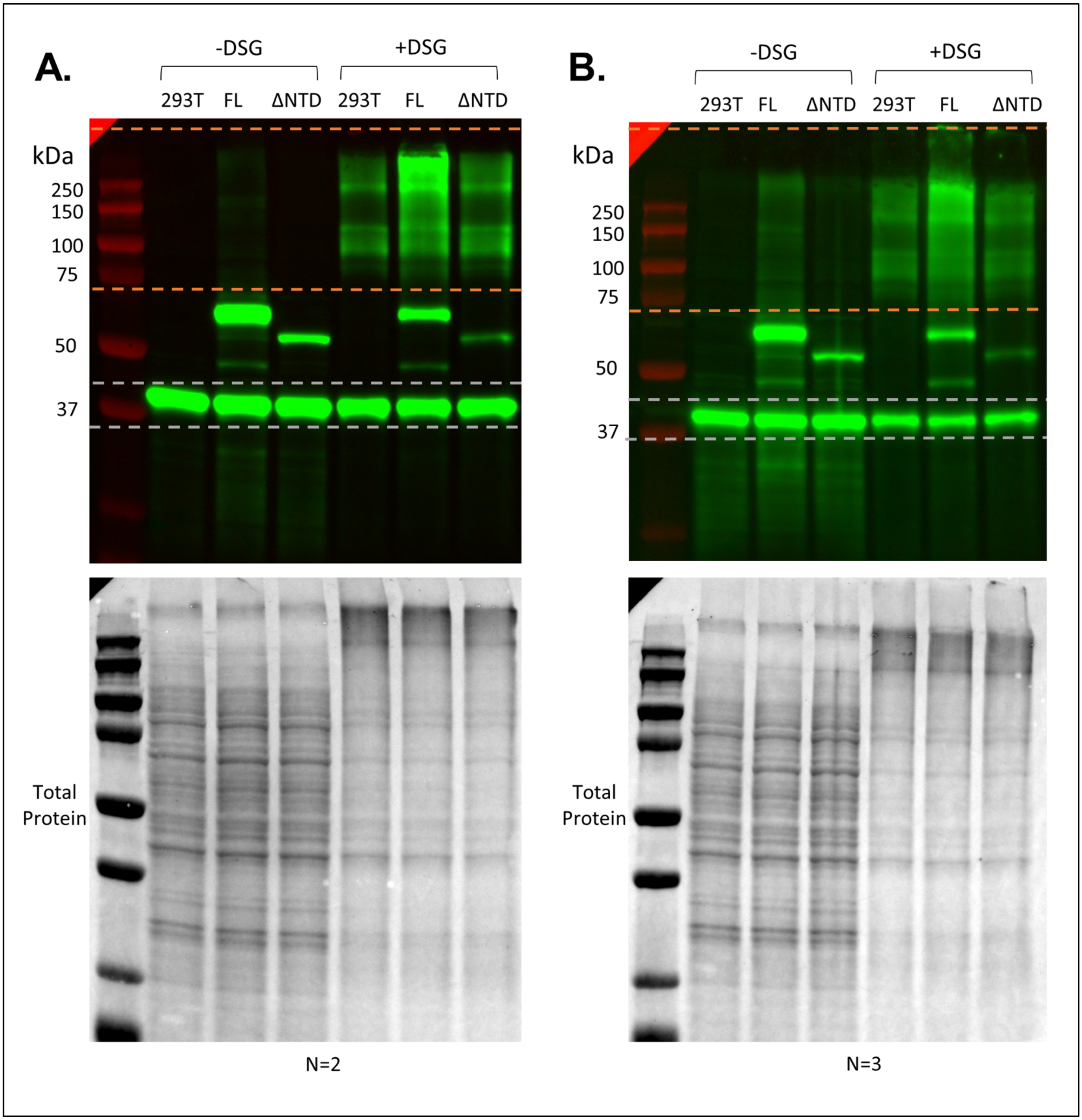


**Supplemental Figure 5. Western blots of un-crosslinked and crosslinked HEK293T cells expressing FL and ΔNTD mNeonGreen biosensors. (A-B)** Each blot shows an independent transfection, harvest and western blotting for TDP-43.

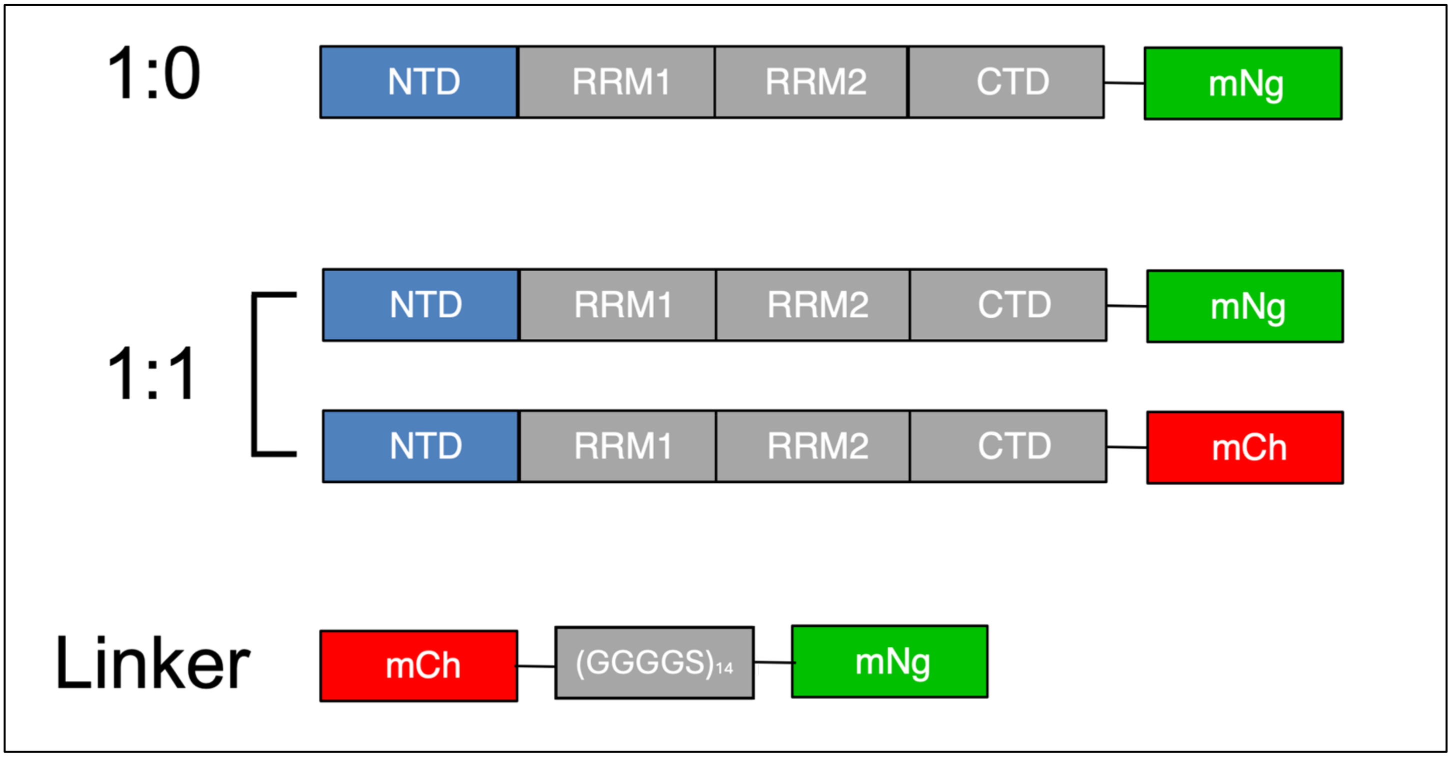


**Supplemental Figure 6. Diagrams of FRET biosensors used in FDA-approved Selleck library screen.**

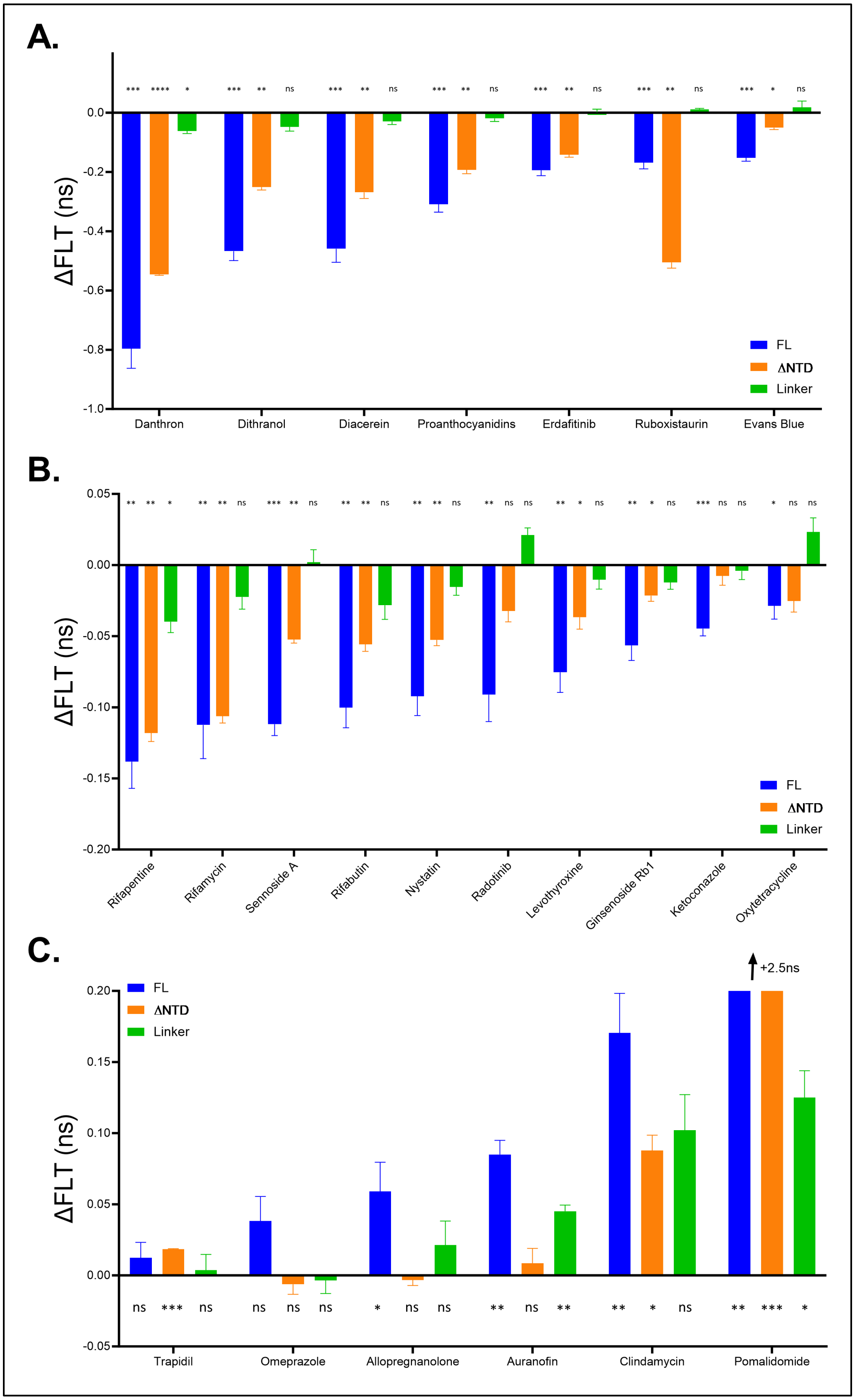


**Supplemental Figure 7. ΔFLT response of all 23 TDP-43 unique hit compounds.** ΔFLT ($FLT_{DMSO}$ - $FLT_{drug}$) for hit compounds organized by effect size and direction (A-C). Blue = FL TDP-43 biosensor, Orange: ΔNTD TDP-43 biosensor, Green: linker control biosensor. Statistics shown are one sample T tests to hypothetical mean of zero (DMSO treatment, * $p < 0.05$, ** $p < 0.01$, ***$p < 0.001$, **** $p < 0.0001$). Data shown are mean ± SEM from N=3 independent experiments.

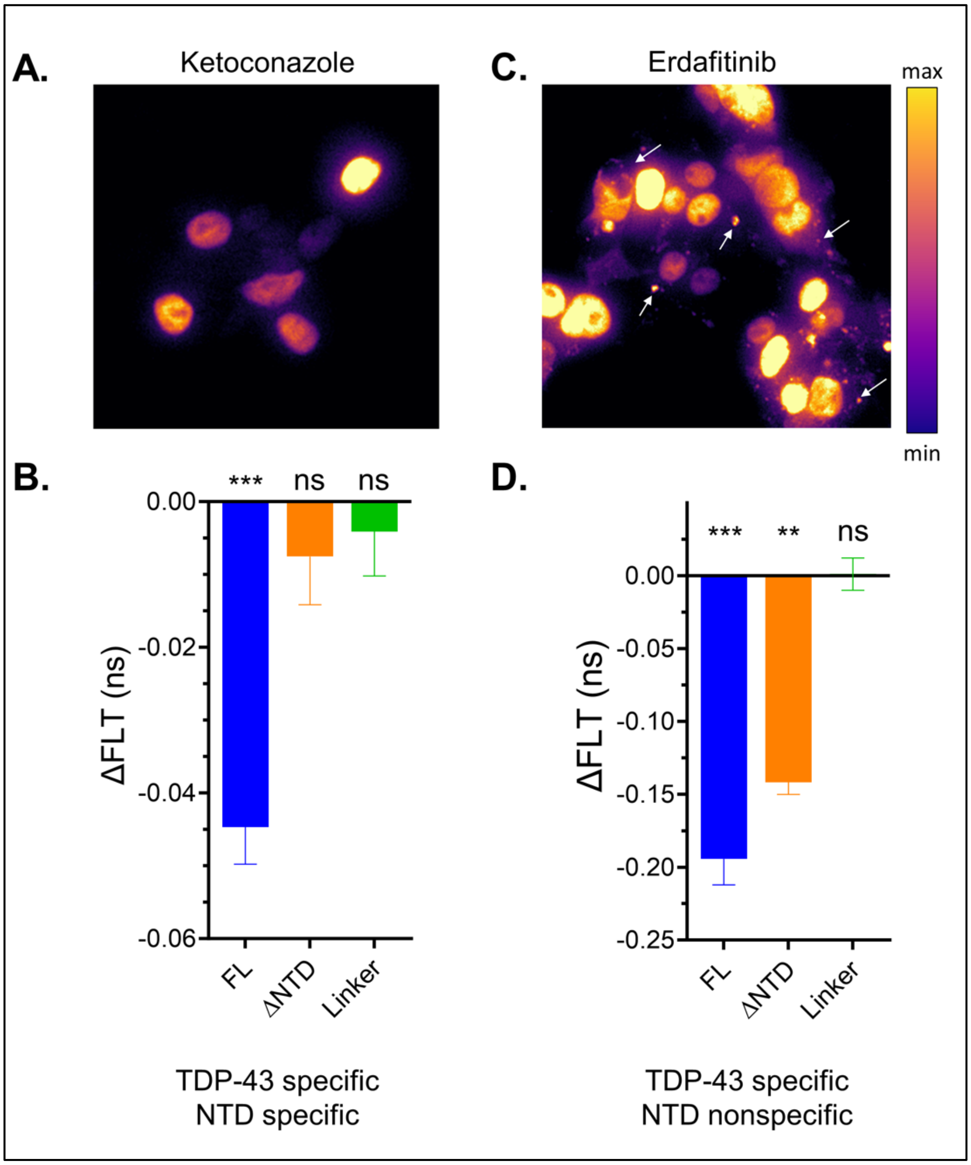


**Supplemental Figure 8. NTD-specific hit ketoconazole does not induce TDP-43 aggregation. (A)** Fluorescence live-cell imaging of TDP-43-mNg expressing HEK293T cells treated with 10 μM ketoconazole for 2 hours. Green fluorescence was mapped to a pseudo-color LUT. **(B)** ΔFLT profile for ketoconazole. **(B)** Fluorescence live-cell imaging of TDP-43-mNg expressing HEK293T cells treated with 10 μM erdafitinib for 2 hours. Green fluorescence was mapped to a pseudo-color LUT. White arrows indicate cytoplasmic TDP-43 puncta induced by erdafitinib. **(B)** ΔFLT profile for erdafitinib. Statistics shown are one sample T tests to hypothetical mean of zero (* $p < 0.05$, ** $p < 0.01$, ***$p < 0.001$). Data shown are mean ± SEM from N=3 independent experiments.

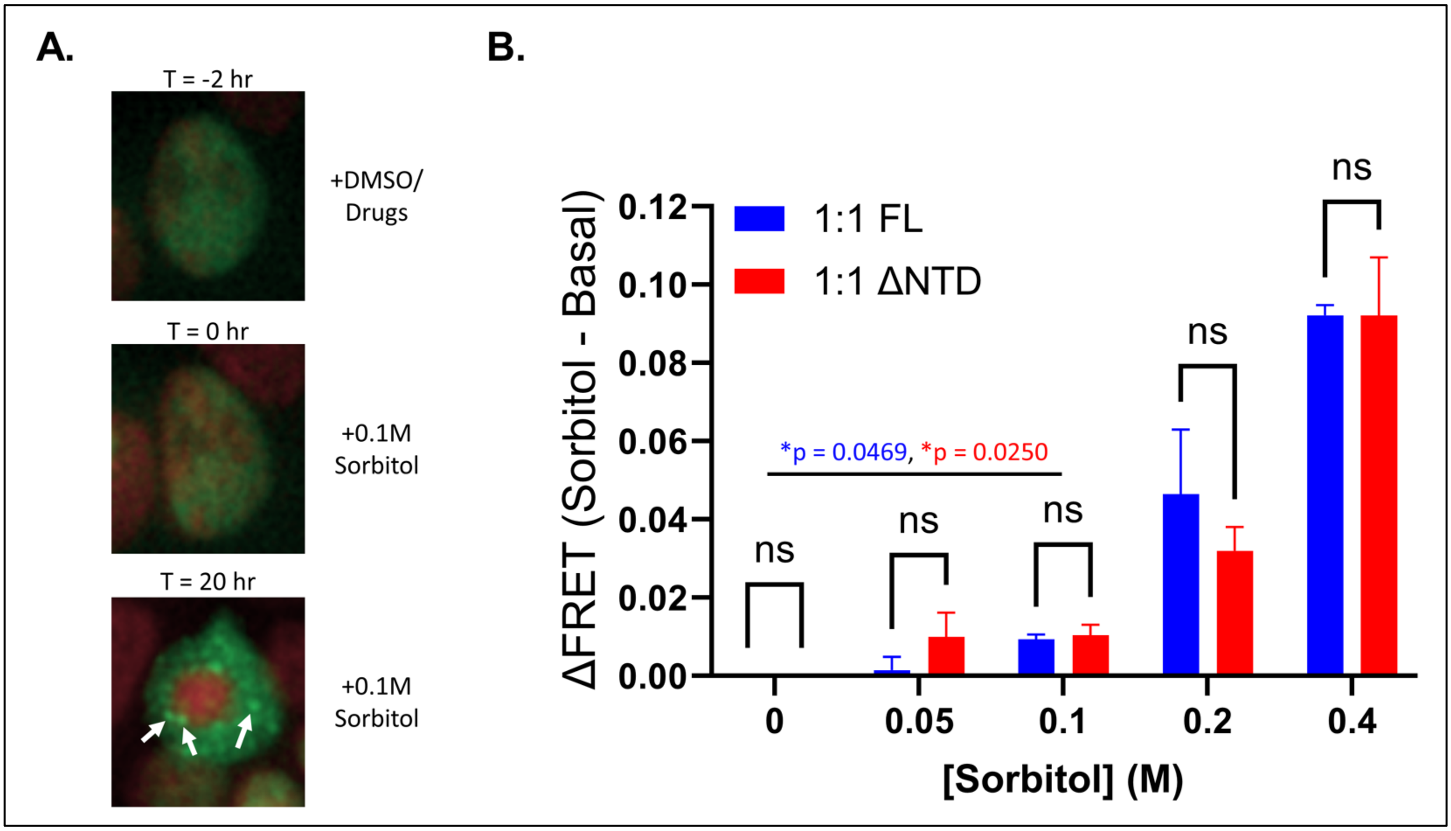


**Supplemental Figure 9. Sorbitol effect on TDP-43 FRET and subcellular localization. (A)** Representative image of TDP43-mNg expressing cell treated with 0.1 M sorbitol for 20 hours. White arrows indicate TDP-43 puncta induced by sorbitol treatment. TDP-43-mNg shown in green and nuclear stain (Hoechst) shown in red pseudo-color. **(B)** Sorbitol titration effect on full-length and ΔNTD TDP-43 biosensor FRET. Statistics shown are two-way ANOVA multiple comparisons with Bonferroni correction (between full-length and ΔNTD bars at single sorbitol concentration) and unpaired T-test (between 0 M and 0.1 M sorbitol for each biosensor, *$p < 0.05$). Data shown are mean ± SEM from N=3 independent experiments.

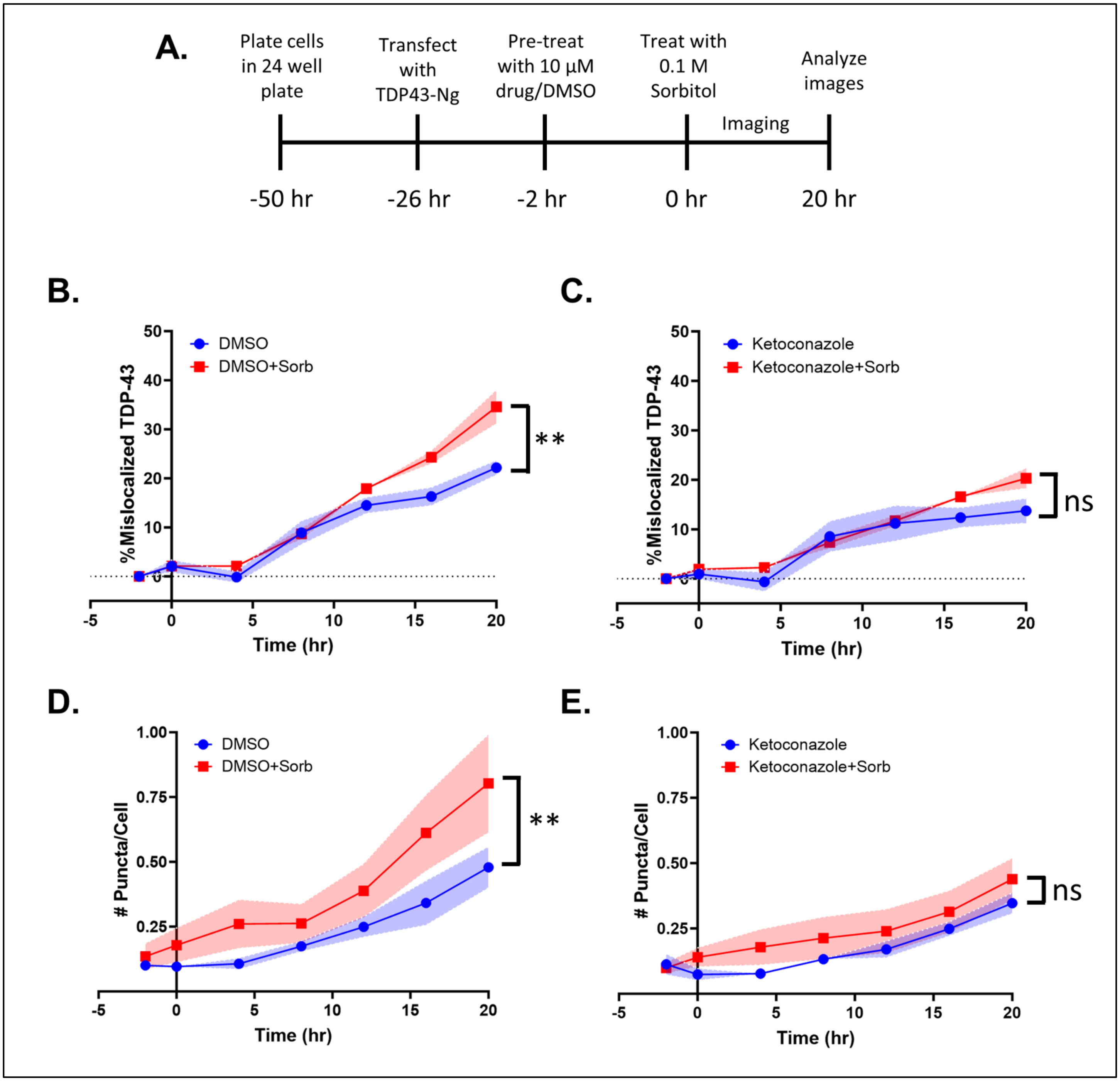


**Supplemental Figure 10. Sorbitol-induced TDP-43 puncta formation and mislocalization experimental design and traces. (A)** Experimental treatment timeline. **(B)** Average TDP-43 mislocalization -/+ sorbitol under DMSO treatment. **(C)** Average TDP-43 mislocalization -/+ sorbitol under ketoconazole treatment. **(D)** Average TDP-43 puncta formation -/+ sorbitol under DMSO treatment. **(E)** Average TDP-43 puncta formation -/+ sorbitol under ketoconazole treatment. Data shown are mean ± SEM from N=3 independent imaging experiments. Statistics shown are two-way ANOVA multiple comparisons with Bonferroni correction (*p < 0.05, **p<0.01, ***p<0.001).

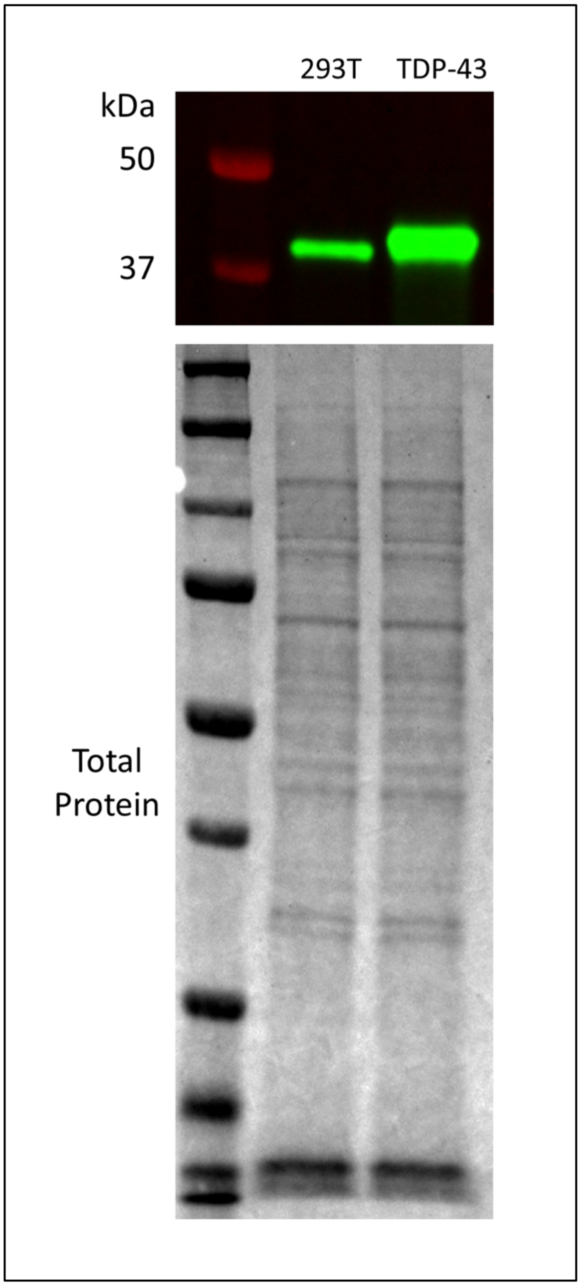


**Supplemental Figure 11. Unlabeled TDP-43 overexpression confirmation in HEK293T cells via western blot.**

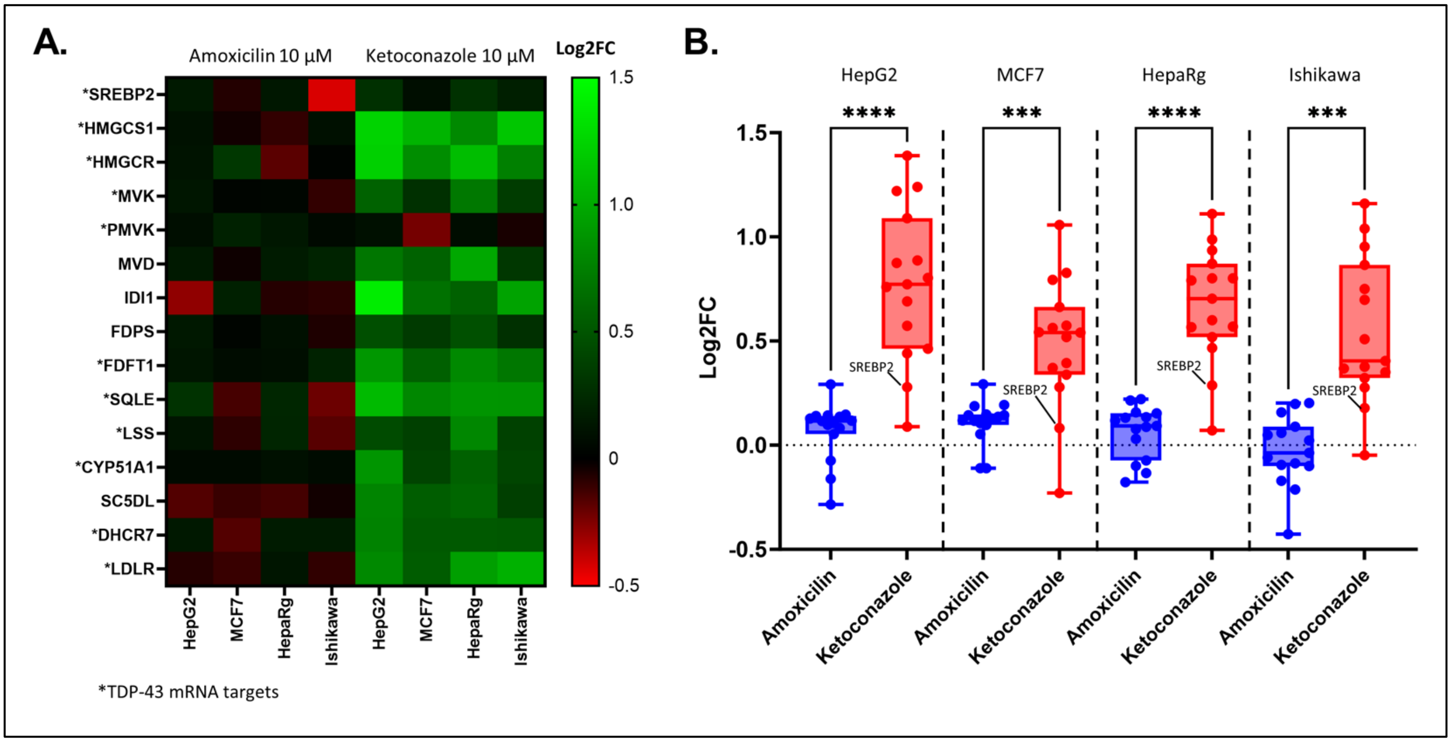


**Supplemental Figure 12. Transcriptomics mining of 10 μM ketoconazole treatment on 4 different immortalized cell lines. (A)** Log2FC (relative to vehicle) heatmap of SREBP2 cholesterol synthesis regulated genes for 10 μM amoxicillin (non-hit in FLT-FRET screen) and ketoconazole (FLT-FRET hit) in HepG2, MCF7, HepaRg and Ishikawa cells. Stars indicate genes known to be mRNA binding targets of TDP-43 protein. **(B)** Summary of Log2FC for genes shown in (A). Statistics shown are one-way ANOVA multiple comparisons with Bonferroni correction (***$p < 0.001$, ****$p < 0.0001$). Data shown are mean from N=3 independent sample treatments.

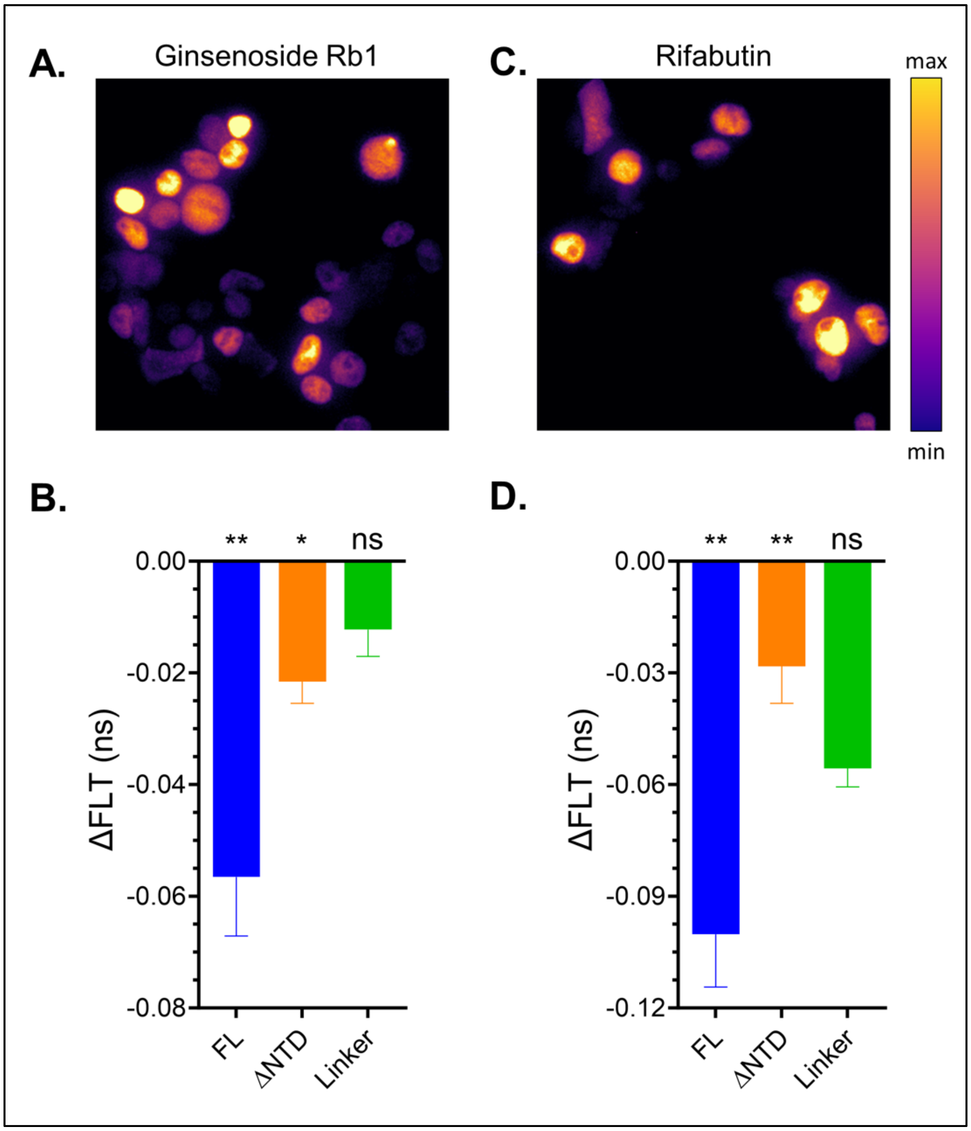


**Supplemental Figure 13. Partial NTD-specific hits do not induce TDP-43 aggregation. (A)** Fluorescence live-cell imaging of TDP-43-mNg expressing HEK293T cells treated with 10 μM ginsenoside Rb1 for 2 hours. Green fluorescence was mapped to a pseudo-color LUT. **(B)** ΔFLT profile for ginsenoside Rb1. **(B)** Fluorescence live-cell imaging of TDP-43-mNg expressing HEK293T cells treated with 10 μM rifabutin for 2 hours. Green fluorescence was mapped to a pseudo-color LUT. **(B)** ΔFLT profile for rifabutin. Statistics shown are one sample T tests to hypothetical mean of zero (DMSO treatment, * $p < 0.05$, ** $p < 0.01$). Data shown are mean ± SEM from N=3 independent experiments.

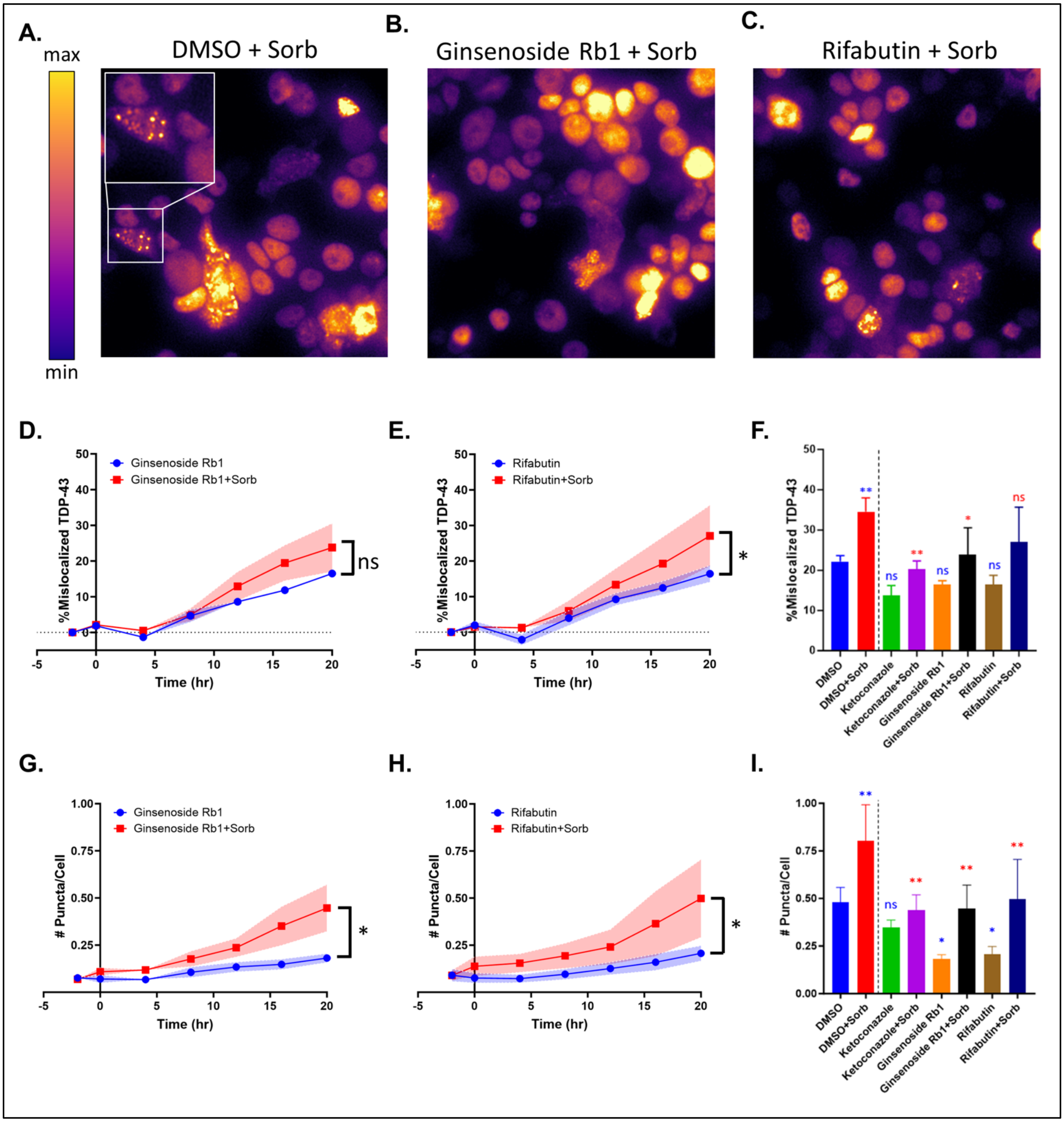


**Supplemental Figure 14. Sorbitol-induced TDP-43 puncta formation and mislocalization under ginsenoside Rb1 and rifabutin treatment.** Fluorescence live cell imaging of HEK293T cells expressing FL TDP-43-mNg treated with 0.1 M sorbitol under **(A)** DMSO, **(B)** 10 μM ginsenoside Rb1 or **(C)** 10 μM rifabutin. Green fluorescence was mapped to a pseudo-color LUT. **(D)** Average TDP-43 mislocalization -/+ sorbitol under ginsenoside Rb1 treatment. **(E)** Average TDP-43 mislocalization -/+ sorbitol under rifabutin treatment. **(F)** Endpoint quantification of TDP-43 mislocalization for all treatments (ketoconazole included for reference). **(G)** Average TDP-43 puncta -/+ sorbitol under ginsenoside Rb1 treatment. **(H)** Average TDP-43 puncta -/+ sorbitol under rifabutin treatment. **(I)** Endpoint quantification of TDP-43 puncta for all treatments (ketoconazole included for reference). Data shown are mean ± SEM from N=3 independent imaging experiments. Statistics shown are two-way ANOVA multiple comparisons with Bonferroni corrections for panels (D), (E), (G) and (H). In panels (F) and (I), blue indicates statistics relative to DMSO-only and red indicates statistics relative to DMSO+Sorbitol (*p < 0.05, **p < 0.01, ***p < 0.001).

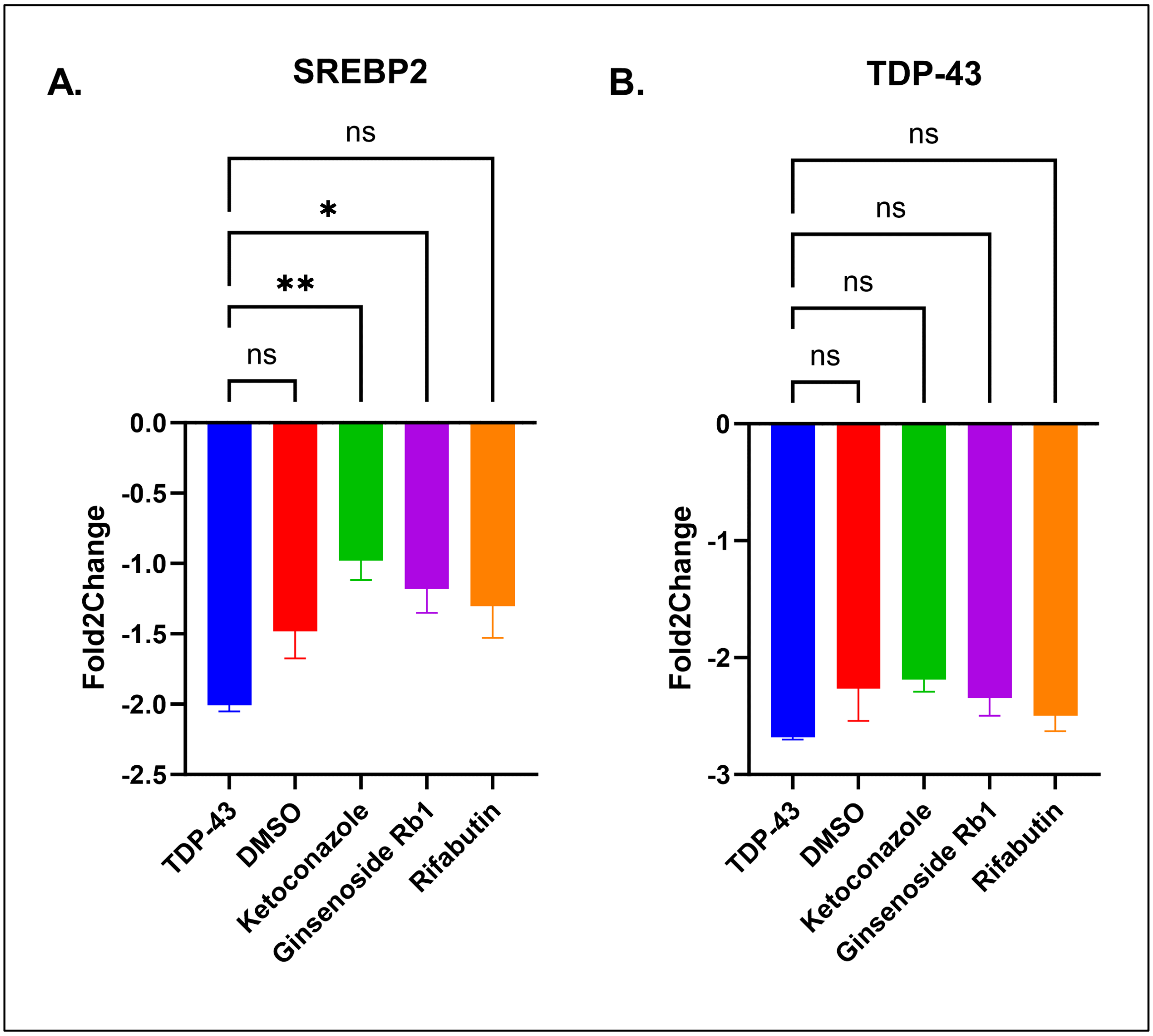


**Supplemental Figure 15. RT-qPCR assay probing for endogenous SREBP2 and TDP-43 under TDP-43 overexpression. (A)** Fold2Change of endogenous TDP-43 mRNA in untreated, DMSO-treated or drug-treated HEK293T cells. **(B)** Fold2Change of endogenous SREBP2 mRNA in untreated, DMSO-treated or drug-treated HEK293T cells. Fold2Changes were calculated relative to untransfected/untreated HEK293T cells and used GAPDH as a housekeeping gene. Statistics shown are one-way ANOVA multiple comparisons with Bonferroni correction against untreated TDP-43 overexpressing cells (blue bar, *p < 0.05, **p < 0.01). Data shown are mean ± SEM from N=3 independent experiments.
